# Spatial Landscape of Pregnancy-Associated Triple Negative Breast Cancer and Mammary Gland Involution

**DOI:** 10.64898/2026.03.09.710650

**Authors:** Darya Veraksa, Kavitha Mukund, David Frankhouser, Lixin Yang, Jerneja Tomsic, Raju Pillai, Jeyasri Venkatasubramani, Daniel Schmolze, Xiao-Cheng Wu, Mary-Anne LeBlanc, Lucio Miele, Augusto Ochoa, Victoria Seewaldt, Shankar Subramaniam

## Abstract

Pregnancy-associated triple negative breast cancer (PA-TNBC) is one of the highest-risk breast cancers, marked by an aggressive phenotype that lacks targeted treatment options. Studies have shown that post-lactational mammary gland involution plays a role in this increased risk. To delineate the underlying mechanisms, our study characterized the transcriptional state of the epithelia and surrounding microenvironment in women with PA-TNBC, comparing those diagnosed pre-involution (PRE) and post-involution (POST, <3 years after delivery). Spatial transcriptomics using the GeoMx Digital Spatial Profiler was performed on treatment-naïve PA-TNBC tissues from 33 women (10 PRE, 23 POST). Regions of interest were segmented with pan-cytokeratin staining. We found that the most prominent transcriptional differences between PRE and POST epithelia occurred in the adjacent non-invasive regions and during the transition into invasive TNBC. POST non-invasive epithelia uniquely showed inflammatory and developmental pathway activation, while the transition into TNBC involved increased chromatin remodeling and cell migration pathways. Further, the tumor microenvironment (TME) in POST showed the highest proportion of immune cells and the highest prevalence of tumor- and immune exhaustion-associated cell states. Finally, a pseudotime analysis of POST transcriptional dynamics found that women diagnosed 1-2 years after delivery exhibited the strongest evidence for inflammatory signaling across the tissue. Our results highlight biological mechanisms distinguishing PRE and POST PA-TNBC across tissue regions and cell types. We emphasize the importance of early detection of malignant molecular signatures in morphologically normal epithelium in post-involution women and suggest that targeting the TME may improve treatment efficacy in post-involution PA-TNBC.

## INTRODUCTION

While triple-negative breast cancer (TNBC) generally carries an unfavorable prognosis^1^, pregnancy-associated TNBC (PA-TNBC) often leads to even poorer clinical outcomes. Historically, pregnancy-associated breast cancer (PABC) has been defined as breast cancer diagnosed during pregnancy or within one year postpartum. Numerous studies have demonstrated that PABC is more aggressive and deadly than non-PABC^2–11^. However, recent large-cohort studies suggest that the heightened aggressiveness of PABC may persist beyond the first postpartum year, proposing to expand the definition of PABC to include women diagnosed up to 5-10 years after delivery^2,3,12^. Interestingly, the decrease in overall survival (OS) in PABC has been attributed specifically to women diagnosed postpartum and not those diagnosed during pregnancy^13^. In support of this, a study of young women’s breast cancer (YWBC; n=619) found that postpartum PABC on its own (<5 years after delivery) had 2.8 times higher risk for metastasis and 2.7 times higher mortality risk than non-PABC cases when adjusting for stage, subtype, and diagnosis year^2^. Further, a meta-analysis of 41 studies found that postpartum PABC (<5 years after delivery) had the worst prognosis when compared with pregnant PABC or PABC overall (OS: HR 1.79; 95 % CI 1.39-2.29)^11^. Thus, although PABC carries a higher risk than non-PABC, evidence points to postpartum PABC, diagnosed within several years after delivery, as the main cause of this disparity.

There is increasing evidence that the aggressive biology of postpartum PABC is driven not by parturition, but rather by mammary gland involution, a two-step process that begins when breastfeeding ceases^14^. Involution causes substantial breast tissue changes as 1) the majority of the secretory mammary epithelial cells undergo rapid death, and 2) the extracellular matrix (ECM) is degraded by matrix metalloproteases, followed by adipocyte differentiation^15^. Previous studies identified epithelial activation of STAT3, TGFβ3, and NFκB signaling as the drivers of this process^16–18^. All three initiate extensive death of alveolar mammary epithelial cells in the first stage of involution; however, STAT3 and NFκB also promote long-term apoptosis resistance through target genes such as CCND1 and BCL2 and have been widely implicated in breast cancer development^16,19^, while TGFβ can induce epithelial-mesenchymal transition (EMT) and promote cancer progression in the long term^17^.

The epithelial changes are accompanied by significant immune infiltration of the mammary gland. As involution proceeds, extensive cytokine signaling promotes a shift from a pro-inflammatory to an immunosuppressive microenvironment enriched with M2-like macrophages, regulatory and Th2 T cells, and fibroblasts^14^. This has led researchers to describe involution as a wound healing-like process and draw parallels with breast tumorigenesis^14,20^. In support of this, mouse model studies have directly linked involution with breast cancer development^14,21,22^. In humans, a study of 16 women with ER+ postpartum PABC identified a transcriptional signature for exhausted T cells and increased cell cycle activity, which was associated with reduced survival in a large validation cohort of YWBC (n=311)^23^. However, the underlying mechanisms are still not fully understood, and the transcriptional state of pre-involution and post-involution PABC has not been well characterized. Furthermore, little is known about PA-TNBC, even though it accounts for 30-40% of all PABC^24^.

To address this critical gap, we assembled a well-defined sample set of 33 women with PA-TNBC (diagnosed during pregnancy or within 3 years postpartum) to uncover deeper mechanistic insights into this disease. The samples were profiled using spatial transcriptomics and whole slide multiplexed imaging to capture the profound impact of mammary gland involution on the breast microenvironment and address the role of tumor-microenvironment interactions in TNBC development. This approach preserved the spatial arrangement of epithelial cells in the context of the surrounding tumor microenvironment (TME) and distinguished between invasive and adjacent non-malignant regions. Our TNBC samples were classified by involution status, rather than as pregnant or postpartum, enabling a precise analysis of how PA-TNBC varies in women who are lactating (pre-involution: PRE) or not (post-involution: POST). To this extent, we first characterized the differences in PRE vs POST across cell populations (epithelia/TME) and tissue regions (non-invasive/invasive). Next, we constructed a pseudo-time trajectory to identify molecular signatures in POST that change as a function of time of diagnosis, measured in days after delivery, aiming to determine the duration of risk following involution. Our results reveal evidence for early malignant changes in non-cancerous epithelial cells and the TME adjacent to TNBC in POST, and an inflammatory signature most strongly apparent in post-involution women diagnosed with PA-TNBC 1-2 years after delivery. These findings can inform effective prevention and therapeutic strategies for this group. To the best of our knowledge, no prior study has examined PA-TNBC with this level of granularity.

## RESULTS

### Study design and ROI selection

PA-TNBC FFPE tissue samples from 33 women (<45 years old) were retrospectively collected through the Louisiana Tumor Registry and annotated for age, stage, race, outcome, and time of diagnosis relative to delivery (Supplementary Table 1). 10 women were identified as pre-involution (TNBC diagnosis during pregnancy or while lactating) and 23 were identified as post-involution (diagnosis after completing involution and within 3 years of delivery). Tissue samples were evaluated by a breast pathologist for lactation-specific morphological features; molecular analysis of lactation was confirmed by elevated expression of lactation-associated genes. 46 slides were acquired and subjected to spatial transcriptomic profiling with GeoMx Digital Spatial Profiler using the GeoMx Human Whole Transcriptomic Atlas (Figure 1a, Supplementary Data, see Methods). 5 of these slides were additionally subjected to spatial proteomic profiling with Orion whole slide multiplexed imaging (see Methods).

**Figure 1:**
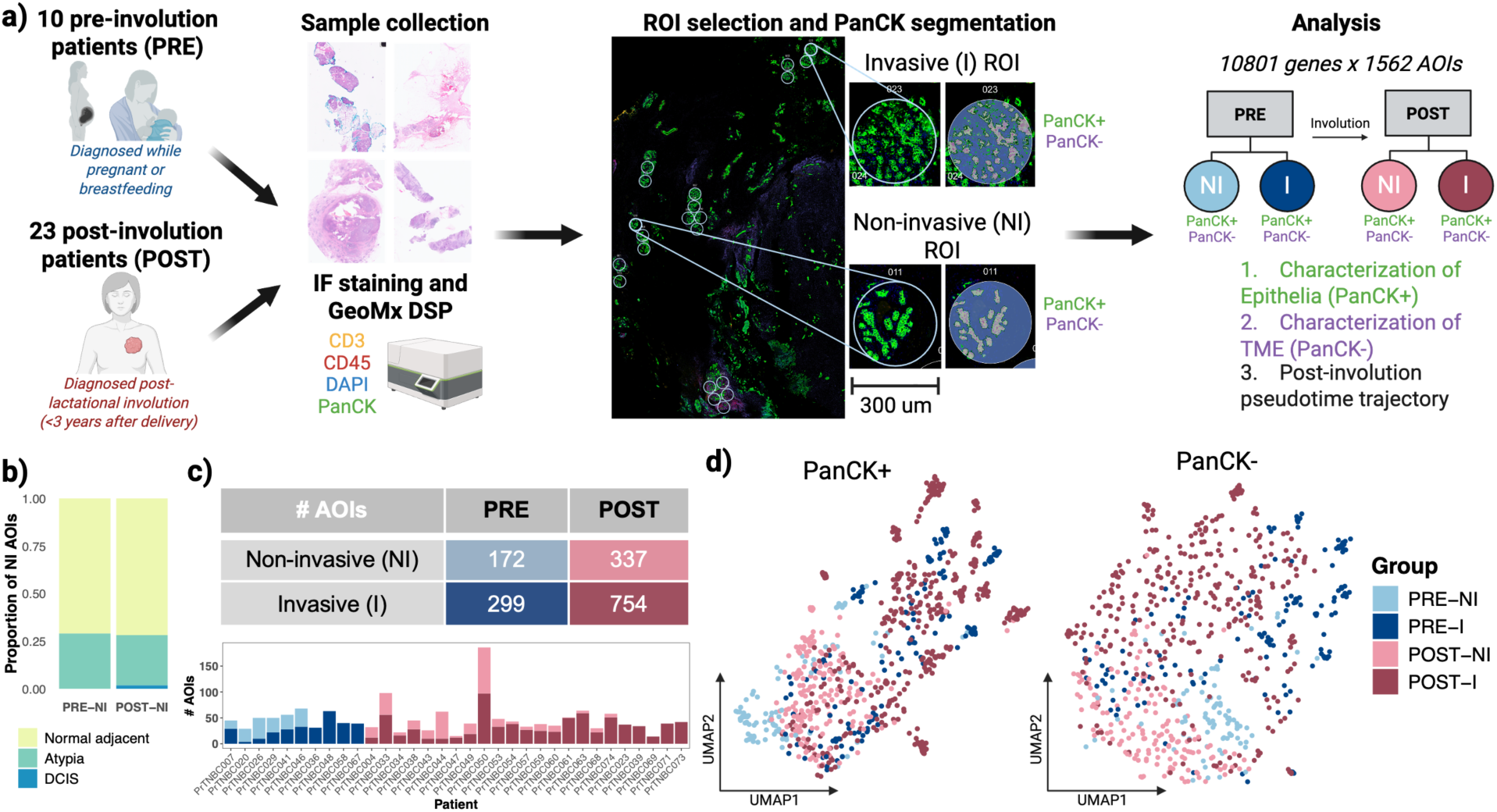
Study design and ROI selection. **a)** Schematic of study design and downstream analysis. PRE: pre-involution; POST: post-involution; NI: non-invasive; I: Invasive; ROI: region of interest; AOI: area of illumination. ROIs were classified by a breast pathologist as containing non-invasive or invasive epithelia. After immunofluorescent (IF) staining, each 300 um ROI was segmented by the green PanCK stain into a PanCK+ AOI (epithelia) and PanCK-AOI (tumor microenvironment - TME). The example shown is part of the tissue excision from PrTNBC004, a post-involution patient. Each AOI was sequenced separately. After quality control the data contained 10801 genes x 1562 AOIs. **b)** Distribution of epithelial types in non-invasive (NI) groups. PRE-NI and POST-NI were mainly comprised of AOIs containing morphologically normal epithelia sampled from tissue adjacent to TNBC (normal adjacent), with some AOIs containing atypia. 3 AOIs with DCIS were in POST-NI. **c)** Table of AOI counts across the four groups being analyzed (PRE-NI, PRE-I, POST-NI, and POST-I) and their distributions across the 33 patients. **d)** UMAP plots of the four groups showed separation between PRE-NI and POST-NI, but not between PRE-I and POST-I in PanCK+ AOIs. PanCK- AOIs showed greater separation between groups.

909 regions of interest (ROIs) were identified across these 46 slides and segmented into 1815 areas of illumination (AOIs) using pan cytokeratin (PanCK) staining to distinguish between PanCK+ epithelia and PanCK- tumor microenvironment. After data processing, 1562 PanCK+/- AOIs derived from 883 ROIs and captured across 10801 genes remained (Supplementary Figure 1a-c, see Methods). Each ROI was annotated for its spatial context (tumor center/tumor edge/isolated foci), and epithelial type (invasive tumor/DCIS/atypia/normal adjacent/distal normal) (see Methods). Invasive (I) epithelia referred to cancerous cells that had spread beyond the basement membrane of a breast duct. Any epithelia not annotated as invasive were referred to as non-invasive (NI) and consisted primarily of morphologically normal ROIs adjacent to tumor tissue, and ROIs that displayed atypical hyperplasia and ductal carcinoma in situ (DCIS) (Figure 1b). These three non-invasive epithelial types exhibited similar expression patterns in preliminary UMAPs (Supplementary Figure 2a). Based on these annotation features, we defined 4 distinct groups of AOIs for downstream analysis: 1) PRE-NI: ROIs from pre-involution TNBC that contained non-invasive epithelia, 2) PRE-I: ROIs from pre-involution TNBC that contained invasive epithelia, 3) POST-NI: ROIs from post-involution TNBC that contained non-invasive epithelia, and 4) POST-I: ROIs from post-involution TNBC that contained invasive epithelia (Figure 1c). Initial UMAPs showed separation between PRE-NI and POST-NI, but not between PRE-I and POST-I in PanCK+ AOIs. PanCK- AOIs showed greater separation between groups (Figure 1d). By systematically comparing these 4 groups across PanCK+ and PanCK- AOIs, we revealed mechanistic insights into tissue state change in PRE and POST PA-TNBC.

### Expression of lactation-associated genes in non-invasive epithelia uniquely identifies involution status

To confirm the classification of each woman as PRE or POST and evaluate the effect of tumor development on lactating epithelia, we analyzed the expression of lactation-associated genes in the PanCK+ AOIs of the two groups. These included the casein proteins (CSN1S1, CSN2, CSN3), genes involved in lactose biosynthesis (LALBA, B4GALT1), fatty acid binding proteins (FABP3, FABP4), and lipogenesis (ACSL1, CD36, LPL). While non-invasive epithelia in PRE highly expressed lactation-associated genes, implying lactation, all non-invasive epithelia in POST suppressed their expression, confirming the pathologist’s annotation of each woman’s involution status (Figure 2a). Further, invasive epithelia in both PRE and POST suppressed the expression of these genes, indicating that cancer formation disrupts normal lactational function. Lactation-associated gene expression in non-invasive epithelia is an agnostic approach to classifying women as pre- or post-involution, which is useful in cases when the histological classification is difficult or the breastfeeding status is unknown.

**Figure 2:**
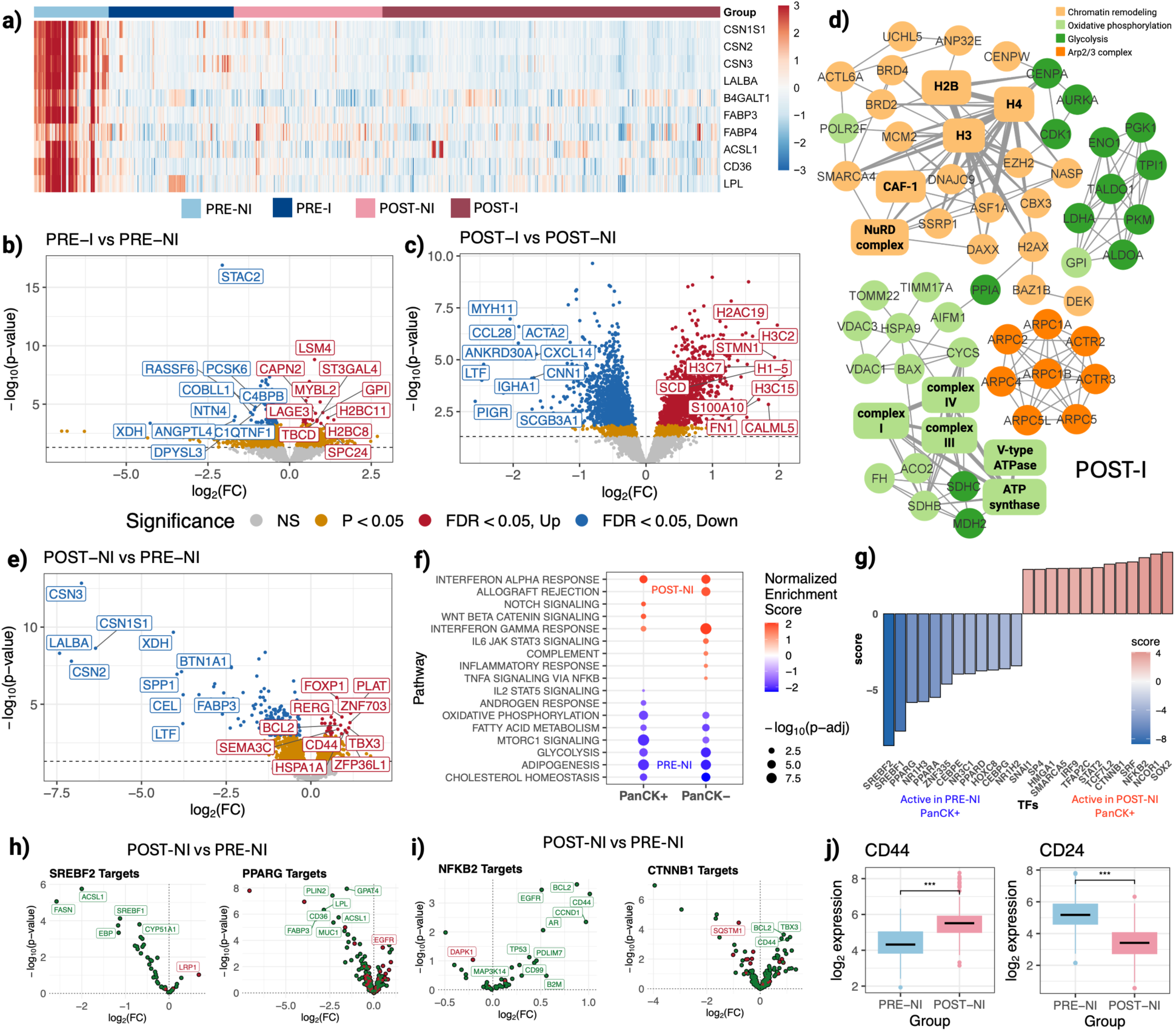
Characterization of epithelia. **a)** Heatmap of lactation-associated genes across all PanCK+ AOIs. Invasive (I) epithelia from pre-involution (PRE) patients and all epithelia from post-involution (POST) patients suppress the expression of these genes, while the surrounding non-invasive (NI) epithelia continue to lactate. Values are row-scaled log2-transformed expression counts. **b)** Volcano plot of PRE-I vs PRE-NI PanCK+ comparison. **c)** Volcano plot of POST-I vs POST-NI PanCK+ comparison. There is a more significant transition in POST than in PRE. **d)** Protein-protein interaction network (confidence > 0.9) of select genes upregulated in POST-I (relative to POST-NI) involved in chromatin remodeling, oxidative phosphorylation, glycolysis, and the Arp2/3 complex. POST-I epithelia are highly transcriptionally active and involved in cell growth and migration. **e)** Volcano plot of the comparison between POST-NI and PRE-NI epithelia shows that lactation-associated genes are differentially expressed in PRE-NI while POST-NI expresses CD44, FOXP1, BCL2, TBX3, and others. **f)** Enrichment of POST-NI vs PRE-NI shows an increase in interferon, Notch, and Wnt-beta catenin pathways in POST. The significant MSigDB pathways found in the epithelial (PanCK+, left) and stromal (PanCK-, right) comparisons are plotted side by side. A positive normalized enrichment score indicates upregulation in POST-NI while a negative score indicates upregulation in PRE-NI. **g)** Transcription factor (TF) target analysis identifies TFs whose targets are differentially active in POST-NI and PRE-NI epithelia. Of note, SREBF1/2, PPARA/D/G, NR1H2/3, NR3C1, and CEBPE/G are differentially active in PRE-NI while SOX2, NCOR1, NFKB2, CTNNB1, TCF7L2, STAT2, IRF9, and SNAI1 are differentially active in POST-NI. **h)** Downstream target genes of SREBF2 and PPARG and their logFC/p-value for the POST-NI vs PRE-NI comparison. Green indicates genes that are activated by the TF and red indicates genes suppressed by the TF. SREBF2 and PPARG targets are mostly overexpressed in PRE-NI (negative log2FC) and include some of the top differentially expressed genes in PRE-NI. **i)** Downstream target genes of NFKB2 and CTNNB1 and their logFC/p-value for the POST-NI vs PRE-NI comparison. Green indicates genes that are activated by the TF and red indicates genes suppressed by the TF. NFKB2 and CTNNB1 targets are mostly overexpressed in POST-NI (positive log2FC) and include some of the top differentially expressed genes in POST-NI. **j)** CD44 expression is significantly higher in POST-NI relative to PRE-NI, while CD24 expression is significantly lower, implying the presence of CD44+/CD24- stem-like cells among POST-NI epithelia.

### The transition from non-invasive to invasive epithelia in post-involution PA-TNBC is marked by activation of chromatin remodeling and cell migration pathways

The comparison of invasive tumors with adjacent non-invasive regions from the same tissue can help identify context-specific molecular changes that underlie tumor progression. To characterize this transition in PRE and POST, we performed differential gene expression analysis between invasive (I) and non-invasive (NI) ROIs. This analysis identified 118 differentially expressed genes (DEGs; FDR < 0.05) when comparing PRE-I vs PRE-NI and 3081 DEGs (FDR < 0.05) in POST-I vs POST-NI (Figure 2b-c, Supplementary Table 2, see Methods). Functional enrichment showed cell cycle activity (downstream of E2F and MYC) and G2-M checkpoint increased in both PRE-I and POST-I, which are common cancer-associated mechanisms responsible for aberrant growth (Supplementary Figure 3a-b, see Methods). POST-I epithelia additionally overexpressed genes involved in metabolism (oxidative metabolism, unfolded protein response, glycolysis, cholesterol homeostasis, and ROS pathways), genes associated with the Arp2/3 protein complex (ARPC1A/1B/2/4/5/5L, ACTR2/3), and 78 genes involved in chromatin remodeling. This indicates a highly active transcriptional state in POST-I epithelia, directed towards cell growth and migration (Figure 2d). Notably, genes with some of the greatest log2-fold changes in the POST-I vs POST-NI comparison (STMN1, FN1, SCD, and KRT81) are known to be associated with TNBC migration^25–28^. In contrast, PRE-I showed weaker enrichment for metabolic pathways, limited to oxidative phosphorylation and unfolded protein response and at a lower significance than POST-I (Supplementary Figure 3a). These results point to dramatic and pronounced epithelial remodeling that occurs in post-involution women during the transition from NI to I, marked by activation of chromatin remodeling and cell migration pathways.

Relative to POST-I epithelia, POST-NI epithelia were enriched for inflammatory signaling pathways: TNFα via NFκB, KRAS, and complement and coagulation signaling (including the complement activators C1QA/B/C, C3, C4B, C7, CFD and the complement inhibitors C4BPB, CLU, and SERPING1) (Supplementary Figure 3a). While the role of NFκB and the innate immune system in involution is well documented^18,29^, the specific role of the complement pathway remains unclear. Notably, both NFκB and complement have been linked to breast cancer progression^19,30^. In contrast, PRE-NI epithelia were enriched for adipogenesis and xenobiotic metabolism, reflecting lipid metabolic activity and detoxification in milk-producing cells (Supplementary Figure 3a). Our results suggest that pro-inflammatory signals persist in the POST-NI epithelia even after the cessation of involution, potentially priming these cells to develop into invasive TNBC through a significant metabolic transcriptional shift.

### Notch, Wnt, and interferon signaling are active in post-involution non-invasive mammary epithelia

Given the differences between PRE and POST in the transition from non-invasive to invasive epithelia, we next characterized the impact of involution on epithelial state, aiming to identify early indicators of tumor aggressiveness. To this extent, we directly compared non-invasive and invasive epithelia between the two groups. We identified 235 DEGs (FDR < 0.05) in the POST-NI vs PRE-NI comparison (Figure 2e, Supplementary Table 2, see Methods). As expected, genes showing increased activity in PRE-NI were associated with pregnancy and lactation, including lactation and butyrophilin family interactions, lipid metabolism, adipogenesis, and cholesterol homeostasis (Figure 2e-f, Supplementary Figure 4). Transcription factor (TF) target analysis on these genes identified SREBF1/2, PPARA/D/G, NR1H2/3, NR3C1, and CEBPE/G as being differentially active in PRE-NI (Figure 2g, see Methods). These TFs regulate lipid biosynthesis and alveolar cell differentiation during milk production and control some of the most highly differentially expressed genes in PRE-NI, such as ACSL1, FASN, LPL, CD36, PLIN2, CSN2, and SPP1 (Figure 2h and Supplementary Figure 5).

In contrast, POST-NI was enriched for interferon, Notch, and Wnt/β-catenin signaling (Figure 2f, Supplementary Figure 4). Notch and Wnt pathways are critical for normal breast epithelial cell differentiation yet have also been implicated as key players in the development of TNBC^31,32^. Further, TF target analysis showed increased activity of SOX2, NCOR1, NFKB2, CTNNB1, TCF7L2, STAT2, and IRF9 in POST-NI (Figure 2g). Notably, NFKB2 activity supports the previous finding that the NFκB pathway is active in POST-NI, while CTNNB1 and TCF7L2 activity support increased canonical Wnt/β-catenin signaling. NFKB2, CTNNB1, and TCF7L2 are known regulators of some of the most highly differentially expressed genes in POST-NI, such as CD44, EGFR, BCL2, TBX3, and SEMA3C, while suppressing downregulated genes in POST-NI, such as CD24 and SQSTM1 (Figure 2i and Supplementary Figure 5). The significant upregulation of CD44 and downregulation of CD24 in POST-NI suggest the stem-ness of these epithelia and an increased ability to form breast tumors^33^ (Figure 2j). This is supported by the significant activity of SOX2, an embryonic transcription factor that is often overexpressed in early basal-like tumors^34^. Further, increased expression of BCL2, an inhibitor of mammary cell death^35^, and FOXP1, a required factor for the activation of quiescent basal mammary stem cells^36^ in POST-NI points to epithelial plasticity and evidence for active mammary cell differentiation and morphogenesis (Figure 2e). Thus, while PRE-NI and POST-NI epithelia both appear morphologically normal, they exhibit different molecular signatures.

Interestingly, we did not find any statistically significant DEGs when directly comparing PRE-I and POST-I epithelia (Supplementary Figure 3c, see Methods). UMAP plots of these AOIs showed that invasive epithelia cluster by sample, while non-invasive epithelia clustered by involution status (Figure 1c and Supplementary Figure 2e). Although the NI-to-I transition differed between PRE and POST, the absence of involution-associated differences among invasive states suggests that, once epithelia acquire an invasive phenotype, they are transcriptionally distinct from NI and independent of involution status. These findings emphasize that the key differences between PRE and POST epithelia emerge from the non-invasive cells, with POST-NI showing elevated interferon, Notch, and Wnt/β-catenin signaling and early signs of epithelial plasticity.

### The tumor microenvironment in post-involution PA-TNBC is inflammatory, mesenchymal, and exhausted

Unlike PanCK+ AOIs that specifically capture epithelial cell expression, PanCK- AOIs capture the transcriptional profiles of a mixture of cell types in the tumor microenvironment (TME). We compared the composition and cell states in PRE and POST PanCK- AOIs to assess the influence of the TME on neighboring epithelia. First, SpatialDecon^37^ characterized the cellular milieu by deconvoluting the proportion of cell types present in each AOI (see Methods). We generated a custom TNBC reference matrix (Supplementary Table 4) using a previously published method from our group^38^, defining expression profiles for 9 cell classes found in the breast TME: plasmablasts, B-cells, myeloid cells (including monocytes and macrophages), CD8+ T cells, CD4+ T cells, other T cell types (including NK cells and cycling T cells), perivascular-like cells (PVL), fibroblast/mesenchymal cells, and endothelial cells. Figure 3a shows the scaled distribution of cell types found in the PanCK- AOIs of the PRE (top) and POST (bottom) slides, arranged by increasing time of diagnosis, measured in days after delivery. There was significantly more immune presence in the TME of POST than in PRE (p<0.05 for B cells, myeloid cells, CD4+/CD8+ T cells, and fibroblast/mesenchymal cells – Figure 3b), which is consistent with the immune alteration that occurs during pregnancy and the immune infiltration expected during the involution process. A recent study comparing PABC diagnosed during gestation vs postpartum also identified an increase in immune cell infiltration in the latter group^39^.

**Figure 3:**
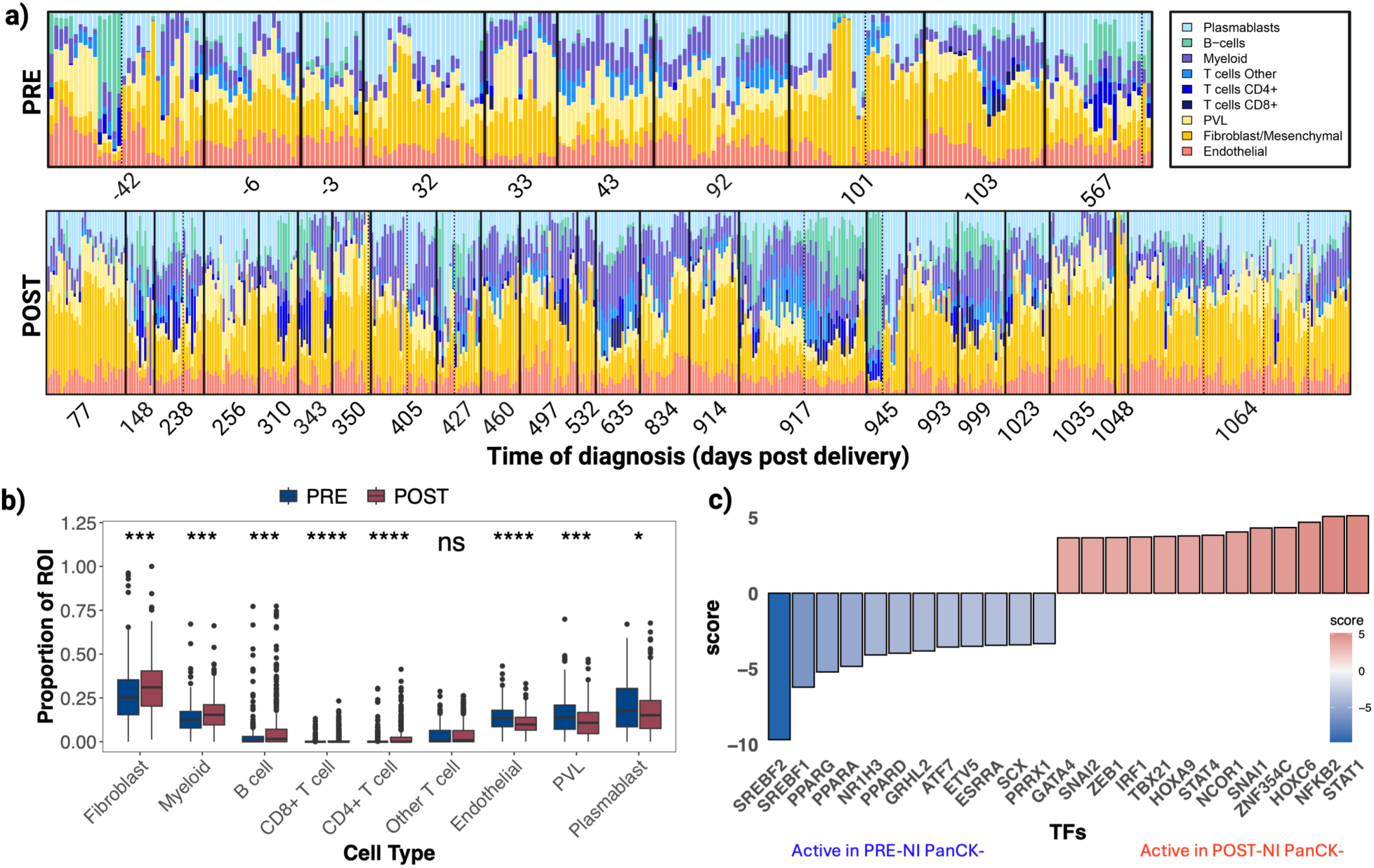
Characterization of TME. **a)** Cell type deconvolution of the tumor microenvironment (TME) in PRE (top) and POST (bottom) PanCK- AOIs using SpatialDecon. Each bar represents 1 AOI. Patients are ordered by their time of diagnosis (measured in days after delivery, with a negative value indicating diagnosis during pregnancy) and separated by a black line. Dashed line distinguishes between multiple slides from the same patient. **b)** Boxplots of the proportions of each cell type across all PRE and POST ROIs indicate significantly more fibroblasts, myeloid cells, B-cells, CD8+ T cells, and CD4+ T cells in POST. There were more endothelial, perivascular-like (PVL), and plasma blast cells in PRE. **c)** Transcription factor (TF) target analysis identifies TFs whose targets are differentially active in POST-NI and PRE-NI TME, many of which are mesenchymal or inflammatory in POST-NI.

The TME surrounding non-invasive epithelia exhibited transcriptional changes aligning with those found in the adjacent epithelia. POST-NI PanCK- AOIs were marked by a strong interferon response, as well as increased IL6/JAK/STAT3, complement, and TNFA via NFKB signaling (Figure 2f, Supplementary Figure 3d-f). Although IL6/JAK signaling activates STAT3, which is necessary for the process of involution, its dysregulation in the microenvironment leads to cancer development and proliferation through an inflammatory feed-forward mechanism^40^. Further, some of the most highly differentially expressed genes in POST-NI PanCK- AOIs were CD44 and BCL2, similar to the epithelia, but also VCAM1, STAT6, and MED28, indicative of an inflammatory and mesenchymal microenvironment (Supplementary Figure 3d, Supplementary Table 3). In support of this, TF target analysis identified STAT1/4, IRF1, NFKB2 (inflammatory) and SNAI1/2, ZEB1, HOXC6 (mesenchymal) as differentially regulating POST-NI TME expression relative to PRE-NI (Figure 3c, see Methods). In contrast, many of the TFs active in PRE-NI TME were the same as those active in PRE-NI epithelia, such as SREBF1/2, PPARA/D/G, and NR1H3, with similar downstream targets (Supplementary Figure 6). This indicates that lactation strongly influences the transcriptional profile of both epithelia and the surrounding TME and is distinctly different from the TME in POST-NI.

To characterize the functional state of the TME in PRE-NI, PRE-I, POST-NI, and POST-I, we employed Ecotyper^41^, a machine learning framework that infers cell state abundances based on carcinoma-derived markers (Supplementary Table 5, Supplementary Figure 8a, see Methods). We observed that the TME exhibited distinct cell states depending on the state of the adjacent epithelia (Figure 4a-b). Endothelial cells, fibroblasts, myeloid cells, and B cells in both PRE and POST were mostly enriched for normal cell states in the TME surrounding NI epithelia, and tumor-associated cell states in the TME surrounding I epithelia. Despite these similarities, POST-I showed the greatest proportion of tumor-associated endothelial cells (top marker genes identified by Ecotyper: ANGPTL2, NID2), CAF1 fibroblasts (POSTN, COL10A1), pro-migratory-like fibroblasts (CA9), and M2 foam cell-like macrophages (AEBP1). These states have been previously associated with shorter survival in a pan-cancer study^41^ and are likely to emerge in a wound-healing-like, post-involution microenvironment. Interestingly, all groups had some form of M2-like myeloid infiltration (M2-like normal enriched (S1PR1) in PRE-NI/POST-NI, and M2-like proliferative (CHI3L2) in PRE-I/POST-I). Using Orion, we showed that ∼80% of all CD63+ macrophages identified across the groups were M2-polarized (CD163+), in tissue regions corresponding to GeoMx ROIs (Figure 4c-d, Supplementary Figure 8b-c, Supplementary Data, see Methods). This suggests that although POST-I is distinguished by higher infiltration of M2 foam cell-like macrophages, all pregnancy-associated tissue exhibits high proportions of M2-like macrophages, albeit with different functional roles.

**Figure 4:**
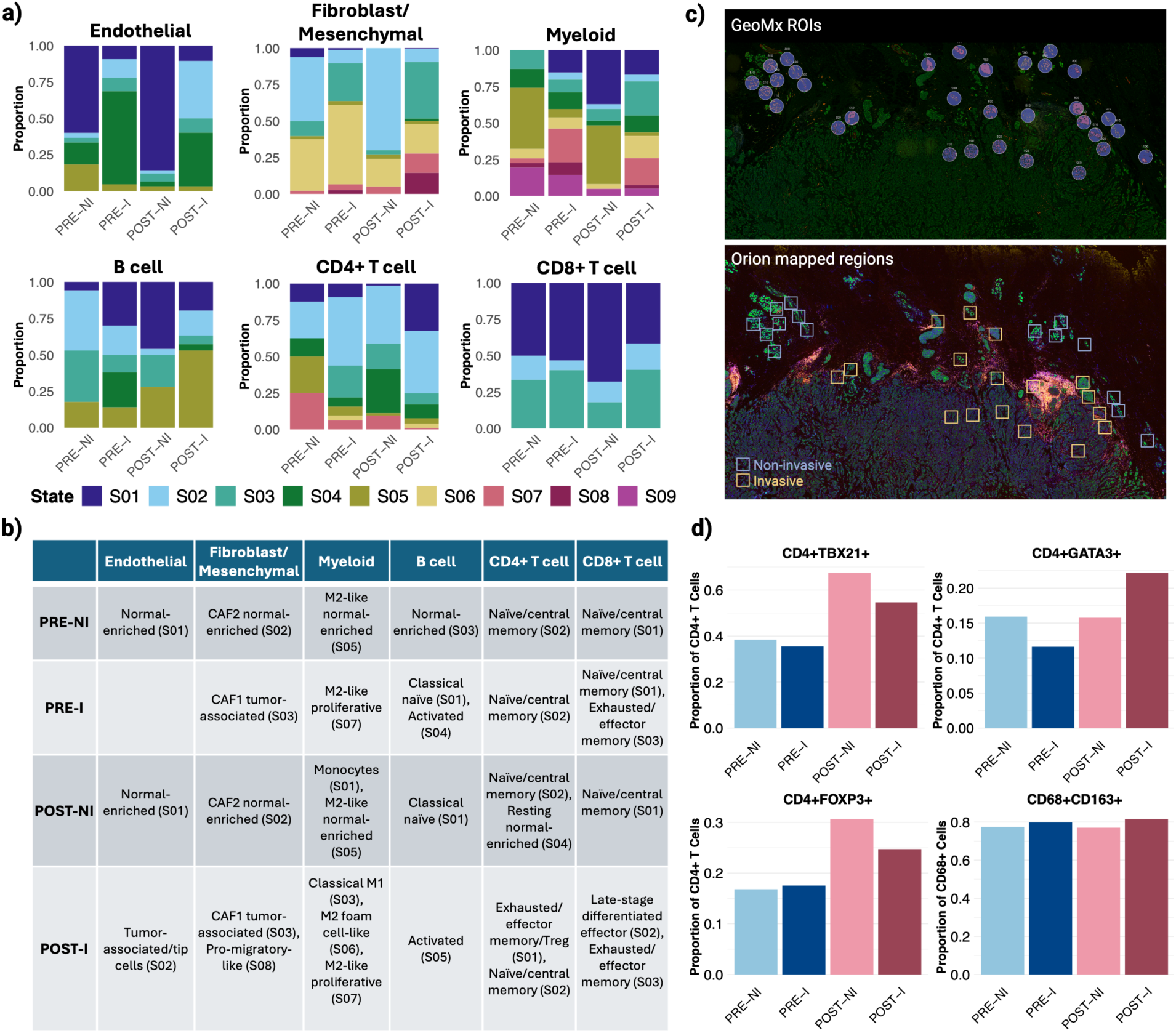
Identifying cell states in the TME. **a)** Cell state deconvolution across PRE-NI, PRE-I, POST-NI, and POST-I using EcoTyper. State labels were taken from the original publication, and full state definitions for each cell type are in Supplementary Figure 8a. **b)** Table of cell states that are most representative of each group. POST-I showed the greatest proportion of tumor-associated/tip endothelial cells, CAF1 fibroblasts, pro-migratory-like fibroblasts, and M2 foam cell-like macrophages. Uniquely, the transition from POST-NI to POST-I showed an increase in exhausted/effector memory/Treg CD4+ T cells and exhausted/effector memory CD8+ T cells. **c)** After pan-immune staining using Orion, 2 PRE and 2 POST slides were annotated with squares (side length: 300 um) to match the location of ROIs sequenced with GeoMx. Additional slide annotations are in Supplementary Figure 8c. Non-invasive/invasive ROI distributions for each slide are in Supplementary Figure 8b. **d)** Barplots of Th1 (CD4+TBX21+), Th2 (CD4+GATA3+), T-reg (CD4+FOXP3+), and M2 macrophage (CD68+CD163+) proportions across the four groups. The greatest proportion of Th1 and T-reg cells were in POST-NI, while the greatest proportion of Th2 cells were in POST-I. All groups exhibited high levels of M2 macrophages.

T cell states were the most distinct in PRE and POST ROIs. Uniquely, the transition from POST-NI to POST-I showed an increase in exhausted/effector memory/Treg CD4+ T cells (CXCR6, CTLA4) and exhausted/effector memory CD8+ T cells (IFNG, GZMB, LAG3), and a decrease in resting (normal-enriched) CD4+ T cells (KLF2) and naïve/central memory CD8+ T cells (BTLA, GZMK) (Figure 4a-b). CD4+ and CD8+ T cell exhaustion has been indicated in immune evasion and subsequent cancer progression^42^, and is consistent with the observation that the TME surrounding POST-I epithelia showed the greatest evidence of immune exhaustion among the four groups. Further, Orion identified a wider variety of T-cell states among groups (Supplementary Figure 9a). Specifically, POST-NI showed the greatest proportion of Th1 (TBX21+) and regulatory (FOXP3+) T cells (out of all CD4 T-cells), while POST-I showed the greatest proportion of Th2 (GATA3+) T-cells. The enrichment of FOXP3+ regulatory T cells and Th1 T cells indicates a mixed immune regulatory and inflammatory environment in non-invasive POST regions, which transitions to a wound-healing-like, immunosuppressive environment nearby invasive TNBC, supporting the Ecotyper results. Overall, these results confirmed that myeloid cells, CD4+/CD8+ T cells, and fibroblast/mesenchymal cells were not only more abundant in the POST-I microenvironment but also existed in an immunosuppressive and exhausted state, creating conditions supporting TNBC progression.

### Inflammatory signaling peaks in post-involution women diagnosed 1-2 years after delivery

Given the key differences in the epithelia and TME between PRE and POST TNBC samples, we next investigated the temporal dynamics in the transcriptional landscape in POST. This could allow us to identify how long after pregnancy a woman remains at risk for aggressive breast cancer due to the altered state of her breast tissue. Based on the distribution of our samples, we binned them into three “pseudo-time” bins: those diagnosed <1 year, 1-2 years, and 2-3 years after delivery (Figure 5a). Next, we performed Sparse Partial Least Squares (SPLS) regression to reduce noise and select key genes for predicting the pseudo-timepoint of an AOI, and ANOVA to assess the ratio of between-sample to within-sample variance (F value) of each of gene identified by SPLS and ensure that no single sample drove the gene patterns (Supplemental Figure 10, see Methods). For each of the four subsets, we removed genes with a high F value (>30, FDR < 0.05). The greatest number of sample-specific genes was removed from the POST-I PanCK+ group (79 genes), which aligns with our prior observation that the expression patterns of AOIs containing invasive epithelia were more sample-specific than any of the other AOI groups. Finally, Mfuzz was utilized for detecting patterns of gene expression across the pseudo-timepoints (Supplementary Figure 10, Supplementary Table 6, see Methods).

**Figure 5:**
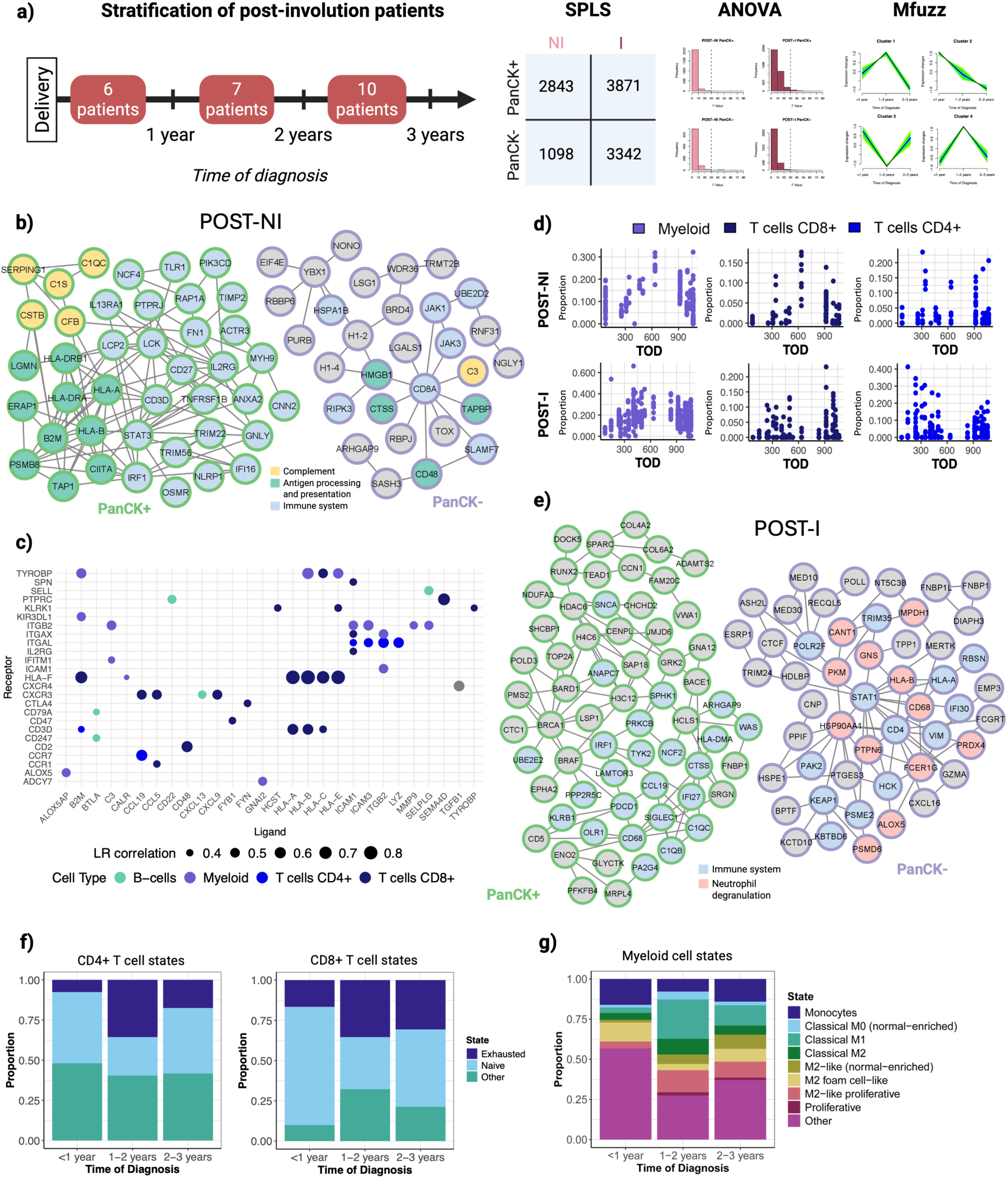
Pseudotime analysis of post-involution patients. **a)** Schematic of the pseudotime analysis in POST patients. After stratifying POST into three groups (those diagnosed <1 year, 1-2 years, and 2-3 years after delivery), we performed SPLS to identify genes in POST-NI PanCK+, POST-I PanCK+, POST-NI PanCK-, and POST-I PanCK-AOIs that were important for separating between the three timepoints. ANOVA removed genes that were too patient-specific and Mfuzz performed fuzzy clustering to identify genes that changed with similar patterns across the three timepoints. **b)** Protein-protein interaction (PPI) networks (confidence > 0.5) of selected genes whose average expression was highest in the non-invasive epithelia (PanCK+, left) and TME (PanCK-, right) of POST patients diagnosed 1-2 years after delivery. Epithelial genes were first reduced to 49 genes associated with the immune system (Reactome); full gene lists are in Supplementary Table 7. There is evidence for activation of JAK/STAT3, an active immune response, and close immune-epithelial interactions in POST-NI ROIs of patients diagnosed 1-2 years after delivery. **c)** Top 50 (by LR correlation) ligand-receptor (LR) interactions in the TME of non-invasive regions in patients diagnosed 1-2 years after delivery, identified using BulkSignalR^43^. The set of LR interactions upregulated in this period were determined by their high gene signature score in Supplementary Figure 9a. The cell type associated with each LR interaction is the highest-scoring (by weight) cell type after performing BulkSignalR cell type assignment. These LR interactions confirm the highly immune-active state of the POST-NI TME in patients diagnosed 1-2 years after delivery. **d)** Distribution of myeloid, CD8+ T cells, and CD4+ T cells across time of diagnosis (TOD) in POST-NI TME (top) and POST-I TME (bottom), measured in days after delivery. Each point represents an AOI. In both POST-NI and POST-I, myeloid proportion appears to peak around 600 days TOD. e) PPI networks (confidence > 0.5) of selected genes whose average expression was highest in the invasive epithelia (PanCK+, left) and TME (PanCK-, right) of POST patients diagnosed 1-2 years after delivery. The invasive epithelia show significant enrichment for immune system-associated genes and surface receptors such as SIGLEC1, KLRB1, and PDCD1 (PD-L1), while the surrounding TME contains CD68, CD4, VIM, and evidence of neutrophil degranulation through genes associated with ficolin-1-rich granule. **f)** Barplots showing the proportions of CD4+ and CD8+ in each time of diagnosis category as determined by EcoTyper. Women diagnosed 1-2 after delivery have the greatest proportion of exhausted CD4+/CD8+ T cells and the lowest proportion of naïve CD4+/CD8+ T cells. **g)** Barplots of myeloid cell state distributions in each time of diagnosis category. The greatest proportion of classical M1, M2-like proliferative, and classical M2 myeloid cells were in the samples diagnosed 1-2 years after delivery.

The greatest inflammatory and immune-associated signaling occurred in POST women diagnosed 1-2 years after delivery (Supplementary Table 7). Specifically, their non-invasive epithelia showed the greatest expression of immune system-associated genes involved in complement and antigen activation and presentation (including MHC class I/II associated genes CIITA and HLA-A/B/DRA/DRB1), as well as the transcription factors STAT3 and IRF1, and FN1 (Figure 5b). The TME near these regions expressed CD8A, TOX, CD48, C3, and JAK1/3, consistent with the enrichment results from POST-NI. These results indicate that the IL6/JAK/STAT3 pathway is likely most active in the women diagnosed 1-2 years after delivery, accompanied by an active immune response and close immune-epithelial interactions through the MHC complex. Further, using BulkSignalR, we inferred ligand-receptor (LR) interactions occurring in the POST-NI TME^43^. Women diagnosed 1-2 years postpartum, even up to 480 days after delivery, demonstrated the greatest evidence for immune activity associated with myeloid, CD4+, and CD8+ T cells (Figure 5c, Supplementary Figure 11, Supplementary Table 8). Interactions involving MHC-associated genes (HLA-A/B/C/E/F), TYROBP, CXCR3, and ITGAL were prominent, indicating a highly active TME in this time period. Interestingly, when examining the proportions of each cell type in an ROI individually as a function of time of diagnosis, we observed significant differences in cell distribution across time, with myeloid and CD8+ T cell proportions peaking in POST-NI around 600 days after delivery (Figure 5d, Supplementary Figure 7). Similarly, the invasive epithelia of these women highly expressed genes involved in the immune system and surface receptors such as SIGLEC1, KLRB1, and PDCD1 (PD-1) (Figure 5e). Upregulated in the surrounding TME were interferon-associated genes, specific immune markers such as CD68 (macrophage) and CD4 (CD4+ T cell), VIM, and ficolin-1-rich granule-associated genes, implying neutrophil degranulation.

Orion performed on 1 representative sample from each pseudo-timepoint confirmed that the sample diagnosed 1-2 years after delivery contained significantly more CD3+ T-cells and CD68+ macrophages per Orion region than the other timepoints (CD3: p < 0.0001, CD68: p<0.05) (Supplementary Figure 9b, Supplementary Data). Furthermore, Ecotyper analysis indicated that the 1-2 year time period had the highest proportion of exhausted CD4+ and CD8+ T cells, classical M1, classical M2, and M2-like proliferative macrophages, and the smallest proportion of naïve T cells (Figure 5f-g). In summary, post-involution women diagnosed 1-2 years after delivery exhibited the greatest evidence for inflammatory and immunomodulatory signaling in both the non-invasive and invasive regions, with the upregulation of IL6/JAK/STAT3 in the non-invasive regions and mesenchymal and exhaustion-associated signaling in the invasive regions. Taken together, these findings indicate that the 1-2 years post-involution period represents a window of heightened biological vulnerability during which risk for aggressive TNBC is elevated.

## DISCUSSION

In this study, we present a comprehensive characterization of pregnancy-associated triple negative breast cancer (PA-TNBC), using GeoMx spatial transcriptomics and Orion whole slide multiplexed imaging to profile invasive and non-invasive regions of breast tissue within their microenvironmental contexts. PA-TNBC is highly aggressive when diagnosed immediately following mammary gland involution, with decreased overall survival and an altered tumor microenvironment (TME)^2–11^. However, the molecular changes occurring during involution that could drive this aggressive phenotype are not fully understood. We evaluate the effect of involution on TNBC development by comparing PA-TNBC from 10 women diagnosed pre-involution (PRE) and 23 women post-involution (POST). Our analysis reveals: 1) transcriptional differences between PRE and POST in adjacent, non-invasive epithelia that may predispose women diagnosed post-involution to develop aggressive invasive cancer, 2) evidence of immunosuppression in the TME near invasive POST epithelia, and 3) peak immune and inflammatory signaling in POST women diagnosed 1–2 years after delivery. Together, these findings suggest that involution-associated changes in non-invasive epithelia and the surrounding TME create a permissive environment for the development of aggressive PA-TNBC.

Given the poor prognosis of TNBC diagnosed after involution, we anticipated that the greatest molecular difference between PRE and POST would be in the invasive (I) mammary epithelial cells. Surprisingly, the most dominant transcriptional difference occurs in the adjacent non-invasive (NI) epithelia (including non-cancerous normal adjacent and atypia), as opposed to the invasive TNBC. PRE-NI epithelia uniquely highly express lactation-associated genes (LALBA and CSN1S1/CSN2/CSN3, among others), suggesting their use as an agnostic marker set for classifying the involutional status of women. In contrast, the state of POST-NI epithelia is characterized by activation of inflammatory pathways through interferon signaling, and developmental pathways through Notch and Wnt/β-catenin. Notably, CD44 and BCL2 expression in POST-NI suggest a population of non-invasive cells that may exist in a plastic and anti-apoptotic state, primed to develop into invasive cancer.

Further, POST-NI epithelia show enrichment for TNFα via NFκB and complement pathways when compared to POST-I, implying that aggressive molecular features may originate prior to tumor development. This early pro-tumorigenic signaling appears to drive a highly significant transition to invasive cancer in POST that is enriched for cell cycle-associated genes and cell migration pathways, a change not seen in PRE. The upregulation of inflammatory pathways, NFκB signaling, Notch, and Wnt/β-catenin has been implicated in promoting epithelial mesenchymal transition (EMT)^44–46^, a process that directs epithelial cells towards a plastic and mesenchymal-like phenotype. Taken together, these findings may reflect early EMT-associated changes emerging in POST-NI epithelia, as seen by elevated CD44 and reduced CD24 expression. The relevance of the contrast between PRE-NI and POST-NI is emphasized by the lack of significantly differentially expressed genes between PRE-I and POST-I (all FDR > 0.05), suggesting that the state of invasive epithelia is less dependent on involution status than the state of non-invasive epithelia. Thus, the most striking differences between PRE and POST epithelia are in the adjacent non-invasive regions, highlighting the critical need for early detection of malignant molecular signatures in morphologically normal-appearing epithelia.

The state of the epithelia is shaped, in part, by the state of the surrounding TME, whose dynamic remodeling during mammary involution has been documented^47^. In our study, the TME surrounding POST-NI epithelia exhibits a signaling profile consistent with that observed in the epithelia. The TME mirrors signatures of complement activation and TNFα via NFκB, while also being enriched for IL6/JAK/STAT3 and targets of mesenchymal transcription factors (SNAI1/2, ZEB1, HOXC6). The elevated fibroblast presence in the POST TME compared to PRE supports this observation and may explain the heightened mesenchymal features observed in POST-NI epithelia. Interestingly, although we do not find significant differences in the invasive epithelia between PRE and POST, we do observe differences in the state of the various cell types in the TME surrounding the invasive epithelia. The POST-I TME exhibits not only the highest proportion of immune cells, but also the highest prevalence of tumor- and immune exhaustion-associated cell states. Of these, the presence of exhausted CD4⁺ and CD8⁺ T cells, which are characterized by the expression of immune checkpoint receptors such as PD-1, CTLA-4, LAG-3, and TIM-3^42^, alludes to a microenvironment that has lost its ability to combat the cancer effectively. In support of this, Orion showed the greatest proportion of GATA3+ CD4+ T cells in POST-I, pointing to a Th2 phenotype which is commonly found in an immunosuppressive microenvironment in breast cancer. Consequently, successful treatments for post-involution TNBC may be more effective by targeting the TME, rather than targeting the invasive cancer. In fact, Tamburini et al. studied the effects of PD-1 blockade in mouse mammary tumor models of postpartum breast cancer and found that it decreased tumor growth during involution and eliminated the exhausted (PD-1+/Lag-3+) CD8+ T cell population^48^.

Our findings indicate that non-invasive epithelia in POST are enriched for inflammatory and developmental pathways, while the surrounding TME is mesenchymal and exhausted. However, the extent of these cell state changes may depend on the interval between delivery and diagnosis. Defining this temporal window is clinically important, as it informs how long women remain at heightened risk of developing aggressive TNBC following pregnancy. To address this, we conducted a pseudo-time trajectory analysis by stratifying our samples into those diagnosed <1 year, 1-2 years, and 2-3 years after delivery. We find that inflammation and T cell exhaustion peak in the post-involution women diagnosed 1-2 years after delivery. Specifically, the non-invasive epithelia and the surrounding TME show signs of coordinated IL6/JAK/STAT3 activity and immune-epithelial interactions through the MHC complex, while the invasive epithelia and surrounding TME express many immune markers as well as PD-L1, indicating an immune presence marked by exhaustion. As expected, the 1- to 2-year time period shows the greatest proportion of exhausted CD4+ and CD8+ T cells, classical M1, classical M2 myeloid, and M2-like proliferative cells, and the smallest proportion of naïve T cells. These cell types are expected in an immunosuppressive involutional microenvironment^14^, yet their presence in the tissue extends past the cessation of involution. We believe this necessitates longitudinal monitoring of women who have recently given birth beyond one year postpartum. Studies extending the post-involution period to 5-10 years after delivery are needed to fully understand how expression patterns change on a longer timescale. Further, although we identified molecular changes across pseudotime, our cohort of 33 women is too small to determine whether this molecular risk affects survival 1-2 years postpartum, highlighting the need for validation in a larger study. Finally, to identify women at risk of developing TNBC prior to onset, a comparison of non-invasive regions adjacent to cancer (what we have studied here) with normal epithelia from post-involution women who remain cancer-free is warranted. Despite these limitations, our work provides a spatially informed molecular characterization of pre- and post-involution TNBC, offering insights that may ultimately guide the development of targeted treatments for women who develop PA-TNBC.

## METHODS

### Sample acquisition

Under the oversight of the Louisiana Cancer Center and the Louisiana State University Institutional Review Board, we identified, from the Louisiana Tumor Registry (LTR) database, we identified 33 pregnancy-associated TNBC cases, 10 women were identified as pre-involution (TNBC diagnosis during pregnancy or while lactating) and 23 were identified as post-involution (diagnosis after completing involution and within 3 years of delivery). Of these 33 women with pregnancy-associated TNBC, 15 women self-identified as White and 16 self-identified as Black. Inclusion criteria for TNBC were: Female breast cancer; ER, PR negative (<1% by immunohistochemistry), absence of HER2-amplification by immunohistochemistry (IHC) and/or fluorescence in situ hybridization (FISH); Age at diagnosis ≥18; Race: Black or White; Any AJCC stage; Documented surgery (lumpectomy or any partial or total mastectomy); documented NACT (Y/N).

### Spatial profiling using GeoMx Digital Spatial Profiler and Orion

Spatial transcriptomic profiling was performed using GeoMx-DSP (Nanostring Technologies). Pre-treatment formalin-fixed paraffin-embedded (FFPE) tissue were cut into 5-mm sections. The first 5 mm section was stained with hematoxylin and eosin (H&E). The second slice was hybridized with the GeoMx Probe Mix for NGS readout (GeoMX Human Whole Transcriptomic Atlas, Cat. #: 121401102, Nanostring Technologies) overnight at 37C. This panel contains 18815 gene probes for 18677 protein-coding genes. Slides were stained with GeoMx Solid Tumor TME Morphology Kit (Cat. #: 121300310, Nanostring Technologies) and GeoMx Nuclear Stain Morphology kit (Cat. #: 121300303, Nanostring Technologies) according to the manufacturer’s protocol. DAPI (blue) was used for nuclei, CD3 (yellow) for T-cells, CD45 (red) for pan-immune cells, and pan cytokeratin (PanCK, green) for epithelial cells. The processed slide was loaded onto GeoMx DSP instrument for ROI selection. A pathologist selected an average of 20 regions of interest (ROIs, diameter = 300 microns) per slide, with replicates representative of different tissue regions (tumor center/edge/isolated foci), epithelial types (invasive tumor/DCIS/atypia/normal adjacent/distal normal), and areas of immune infiltration. The ROIs were segmented into PanCK+ (epithelial) and PanCK- (tumor microenvironment - TME) areas of illumination (AOIs) and individually sequenced. Probes were collected and transferred to a PCR plate for library prep using Seq Code primers (GeoMx Seq Code Pack, Cat. #:121400205-121400206, Nanostring Technologies). Libraries from each AOI were pooled, purified by AMPure XP beads (A63880 Beckman Coulter), and resuspended in a volume of elution buffer proportional to the number of pooled AOIs. Libraries were assessed using an Agilent TapeStation, diluted to 1.6 pmol/L, and sequenced (paired-end 2×27) on Illumina NextSeq2000, with a coverage of 100 reads. FASTQ files were converted into DCC files by GeoMxNGSPipeline software and utilized for all downstream processing. A total of 1815 AOIs derived from 909 ROIs were captured across 46 slides.

Spatial proteomic profiling was performed using Orion whole slide multiplexed imaging. Consecutive tissue sections were cut from 5 (2 PRE and 3 POST samples) of the same tissue blocks profiled by GeoMx and processed using the Orion multiplex immunofluorescence protocol (Akoya Biosciences). The markers measured included: Pan-CK (ArgoFluor 874), CD3e (ArgoFluor 548), CD4 (ArgoFluor 760), CD8a (ArgoFluor 624), PD-1 (ArgoFluor 782), FOXP3 (ArgoFluor 724), TBX21 (ArgoFluor 572), GATA3 (ArgoFluor 676), GrzB (ArgoFluor 520), and CD45RO (ArgoFluor 706).

### Data visualization and processing

All data was processed using R (version 4.5.0) and visualized with ggplot2 (3.5.2) and Biorender. *Set.seed(42)* was used throughout the analysis for umap (0.2.10.0), SPLS (2.3-2), Mfuzz (2.68.0), and BulkSignalR (1.0.4).

### GeoMx data quality control and preprocessing

Quality control (QC) and data preprocessing were performed using GeomxTools (3.11.0) and NanoStringNCTools (1.15.0) in accordance with developer recommendations. Supplementary Table contains the custom annotation file. Probe assay metadata were from Hs_R_NGS_WTA_v1.0.pkc. After pooling all AOIs, the following QC flags were used: *minAOIReads = 1000, percentTrimmed = 80, percentStitched = 80, percentAligned = 80, percentSaturation = 50, minNegativeCount = 1, maxNTCCount = 5000, minNuclei = 30, minArea = 1000, LOQ = 2*. AOIs with less than 5% of genes detected and genes not seen in at least 10% of the AOIs were excluded. After QC there remained 10801 genes across 1562 AOIs (from 883 ROIs). The raw counts were Q3-normalized and log2-scaled.

### Differential analysis using linear mixed models

All differential analysis was performed using linear mixed models (LMM) from lme4^49^ (1.1-37). LMMs are linear models that incorporate random effects to account for the fact that multiple ROIs are sampled from the same tissue. Because some women (samples) were sampled across multiple slides, a random intercept was included in the form *sample/slide*, as is appropriate for the nested structure of the data. We performed two types of comparisons: 1) For *within-slide* comparisons between features co-existing on a slide (such as invasive vs non-invasive epithelia in PRE), we used the following formula: *log2(gene) ∼ invasive_status + (1 + invasive_status | sample/slide)*. Slides without both features of interest were excluded. 2) For *across-slide* comparisons between the same feature in different groups (such as invasive epithelia in PRE vs invasive epithelia in POST), we used the following formula: *log2(gene) ∼ involution_status + (1 | sample/slide)*. Differentially expressed genes (DEGs) with FDR < 0.05 were considered significant.

### Enrichment and protein-protein interaction networks

We performed enrichment using fGSEA^50^ (1.34.2) and MSigDB Hallmark^51^ pathways. Genes were ranked by *sign(logFC)*-log10(p-value)* and only pathways with p-adj < 0.05 were displayed. Protein-protein interaction (PPI) networks were visualized on Cytoscape^52^ (3.9.1) using a background downloaded from StringDB^53^ (9606 v12.0). The interactions were filtered for interaction score > 500, and duplicated edges and self-loops were removed. The final source network had 19220 nodes and 569217 edges. The POST-I PanCK+ PPI (Figure 2d) was further filtered to high confidence (interaction score > 900) and included significant (FDR < 0.05) genes found to be involved in oxidative metabolism and glycolysis (MSigDB), the Arp2/3 complex, and chromatin remodeling (GOBP). Genes involved in H2B, H3, and H4 histone families, ATP synthase, V-type ATPase, NADH:ubiquinone oxidoreductase (mitochondrial complex I), cytochrome bc1 (mitochondrial complex III), cytochrome c oxidase (mitochondrial complex IV), nucleosome remodeling and histone deacetylase (NuRD) complex, and chromatin assembly factor 1 (CAF-1) complex were grouped together for simplicity.

### Transcription factor target analysis

Transcription factor (TF) activity inference was performed for each comparison using DecoupleR^54^ (2.14.0), using CollecTRI (43159 TF-target interactions) as the background. A statistic including the direction of change and significance was calculated for each gene with *sign(logFC)*-log10(p-value),* and a univariate linear model was constructed to infer differential TF activity from these values. The top 25 scoring TFs were displayed.

### Cell-type deconvolution of the TME

SpatialDecon^37^ (1.18.0) was used to deconvolute the cell types of the tumor microenvironment (TME) in each PanCK- AOI. A previously described method^38^ was used to generate a custom TNBC-specific reference matrix instead of the default SafeTME matrix, which is largely derived from peripheral blood mononuclear cells and may not accurately represent the breast microenvironment. scRNAseq data from the TNBC microenvironment was downloaded (https://singlecell.broadinstitute.org/single_cell/study/SCP1039/a-single-cell-and-spatially-resolved-atlas-of-human-breast-cancers#study-download)^55^ and processed with Seurat^56^ (5.2.0) to obtain 11937 features x 25188 cells (B cells: 1839, Endothelial: 900, Fibroblast/Mesenchymal: 1946, Myeloid: 5168, Plasmablasts: 853, PVL: 748, CD4+ T cells: 6614, CD8+ T cells: 5137, Other T cells: 1983). The features were further reduced to 1069 breast-specific genes^38^. *create_profile_matrix()* was used to generate a matrix containing 1069 genes across 9 cell classes: plasmablasts, B-cells, myeloid cells (including monocytes and macrophages), CD8+ T cells, CD4+ T cells, Other T cell types (including NK cells and cycling T cells), perivascular-like cells (PVL), fibroblast/mesenchymal cells, and endothelial cells (Supplementary Table 4). After calling *runspatialdecon(),* the predicted contributions of each cell type in each AOI were scaled to 1 and plotted. A Wilcoxon test was used to compare proportion distributions between PRE and POST for each cell type and identify those that are significantly differential between the groups.

### Identifying cell states in the TME

Ecotyper^41^ was used to identify the cell states of each deconvoluted cell type. The raw expression data (post-probe QC but prior to Q3 normalization) were converted to TPM and inputted to the *EcoTyper_recovery_bulk.R* script that recovers previously-identified cell states in carcinomas. In this case, each AOI is treated as a mini-bulk sample because it contains a mixture of cell types. As previously described^38^, we multiplied the state abundances returned by Ecotyper with the cell type proportions returned by SpatialDecon to obtain an accurately scaled state abundance estimate for each AOI. Lastly, AOIs were designated as containing cells of each cell type in a particular cell state if their calculated state abundance exceeded the 80^th^ quantile of the distribution of state abundances for each cell type. The final state assignments are in Supplementary Table 5. The resulting state assignments were scaled to 1 and plotted for each of the overlapping cell types between Ecotyper and SpatialDecon (endothelial cells, fibroblasts, myeloid cells, B cells, CD4+ T cells, and CD8+ T cells). The description of each cell state was taken from Supplementary Table 4 of the Ecotyper publication. To make the boxplots in Figure 5e, all PanCK- AOIs with nonzero cell presence (by SpatialDecon) of either CD4+ T cells, CD8+ T cells, or myeloid cells were considered for each time point.

### Orion image analysis

The Orion images were processed and analyzed using QuPath v0.5.1. Square regions (side length: 300um) were manually annotated on each image to match the size and location of ROIs sequenced with GeoMx and labeled as invasive or non-invasive. Cell segmentation was performed using *Cell Detection* with default settings, and intensity features were calculated using *Add Intensity Features* for each biomarker stain channel. Using the mean intensity from each cell, a cell positivity threshold was determined for each channel by fitting the observed intensities with Gaussian-mixture model assuming two distributions. The point of intersection between the means of the two detected distribution was used as the positivity threshold, and all cells with mean intensity above this value were labeled as positive for each biomarker. 2 PRE and 2 POST samples (selected by the slides with the most even NI/I ROI distributions) were used for the comparative analysis described in Figure 4c-d, while the 3 POST samples (1 representing each time of diagnosis category) were used for the quantification of CD3+/PanCK+ cells across pseudotime. For the comparative analysis, the number of cells identified across square areas in NI/I regions from each patient are shown in Supplementary Figure 8b. An upset plot was created to visualize all unique T cell subtypes (with >50 cells) based on combinations of markers among CD3+ cells (Supplementary Figure 9a). CD4+ T cells were identified as CD3+CD4+CD8-, and proportions of CD4+ T cells that were Th1 (CD3+CD4+CD8-TBX21+), Th2 (CD3+CD4+CD8-GATA3+), and T-regulatory (CD3+CD4+CD8-FOXP3+) were calculated to create the bar plot in Figure 4d. Macrophages were identified as CD68+, while M2 macrophages were CD68+CD163+. To confirm T cell presence across pseudotime, 1 representative patient from each timepoint (POST diagnosed <1 year, 1-2 years, and 2-3 years) was selected for Orion staining. CD3+, PanCK+, and CD68+ cells were quantified and compared in Supplementary Figure 9b using the Wilcoxon test.

### Pseudotime analysis in the post-involution women

The post-involution women were binned into three categories based on their time of diagnosis (TOD, measured in days past delivery): those diagnosed <1 year, 1-2 years, and 2-3 years after giving birth. Sparse Partial Least Squares (SPLS) regression was performed separately on log-normalized counts in POST-NI PanCK+ (181 AOIs), POST-NI PanCK- (156 AOIs), POST-I PanCK+ (411 AOIs), and POST-I PanCK- (343 AOIs) using spls (2.3-2). SPLS is well-suited for datasets with many more features (genes) than samples (AOIs) because it reduces noise and extracts only the necessary genes for predicting the pseudo-timepoint of an AOI^57^. Prior to running *spls()*, 10-fold cross-validation with *cv.spls()* was run to identify the optimal sparsity tuning parameter (eta) and number of latent components (K) for each group of AOIs, listed in Supplementary Table 6. SPLS identified 2843 features as important in POST-NI PanCK+, 1098 in POST-NI PanCK-, 3871 in POST-I PanCK+, and 3342 in POST-I PanCK-. Next, ANOVA was performed to assess the ratio of between-sample to within-sample variance (F value) of the expression of each gene returned by SPLS, with the formula *value ∼ sample* (Supplementary Figure 8). Genes with F-value > 30 (FDR < 0.05) were removed from the gene lists returned by SPLS. There remained 2839 genes in the POST-NI PanCK+ AOIs, 1092 genes in the POST-NI PanCK- AOIs, 3792 genes in the POST-I PanCK+ AOIs, and 3327 genes in the POST-I PanCK-AOIs that were important for predicting the TOD category.

Soft fuzzy clustering on each gene set was performed using Mfuzz^58^ (2.68.0)*. mestimate()* and *Dmin()* were used to identify the optimal fuzzifier parameter and number of clusters, respectively (Supplementary Table 6). *Dmin()* calculates the minimum centroid distance between two cluster centers, and its value decreases slower after reaching an optimal number of clusters (Supplementary Figure 8). For consistency we chose c=10 clusters for each gene set. The mean expression of each gene per TOD category was calculated, genes were standardized to have mean of 0 and standard deviation of 1 across the three categories, and clustering was performed using *mfuzz()* (Supplementary Figure 8). The clusters were evaluated for each gene set and genes coming from clusters with similar trends (ie increasing over time, decreasing over time, peaking at time 2, etc) were combined and evaluated for enrichment and protein-protein interactions. Genes that peaked at time 2 (diagnosis 1-2 years after delivery) are listed in Supplementary Table 7.

### BulkSignalR for identifying ligand-receptor interactions

We used BulkSignalR^43^ (1.0.4) to infer ligand-receptor (LR) interactions in the PanCK- AOIs in invasive (343 AOIs) and non-invasive (156 AOIs) regions. Each PanCK- AOI contains more than one cell, so it is appropriate to treat it as a mini-bulk sample. BulkSignalR infers LR interactions that are statistically likely to be present in each sample by calculating correlations between L and R, and R and its downstream-regulated target genes (T). It identifies LR and RT null distributions of Spearman correlations by repeatedly randomizing the input data, then tests the significance of each LR correlation in the samples under the null. Q3-normalized expression data (prior to log2 scaling) were used to create two BulkSignalR objects for all POST-NI PanCK- AOIs and POST-I PanCK- AOIs separately, using *min.count = 0, prop = 0, normalize = FALSE, method = ‘UQ’*. *BSRinference()* was calculated with *min.cor = 0.25*, and after reducing to the best pathway only those with *q < 0.05* were retained. LR interactions were correlated with the cell type proportions identified by SpatialDecon to infer the cell types in each AOI that were most likely associated with each LR interaction (*q < 0.05* and *weight > 0.25*). All results are in Supplementary Table 8. Significant LR interactions in the two BSR objects were assigned a gene signature score based on a weighted sum of ligand, receptor, and target gene z-scores and plotted in order of time of diagnosis (Supplementary Figure 9).

## Data and Code Availability

DCC files, the processed GeoMx data object, and the Orion data object can be found under the accession GEOXXXXXXX.

## ACKNOWLEDGEMENTS

The authors thank Dr. Jason Swedlow and Dr. Josh Titlow from the Wellcome Leap Foundation for their support and scientific guidance.

## AUTHOR CONTRIBUTIONS

D.V designed and performed the computational analyses, with assistance from K.M in design and J.V in figure generation. D.F processed the GeoMx slide images, and performed square region annotation, cell identification, and fluorescence quantification of Orion data. L.Y, J.T, and R.P designed the GeoMx sequencing assays and generated the data. D.S and V.S. were the lead pathologists in the study and identified all GeoMx ROIs considered in the study. X-C.W, M-A.L, L.M and A.O selected all samples and associated metadata from the LTR database. S.S and V.S designed and supervised the study. D.V wrote the first draft of the manuscript, with revisions by K.M, V.S, and S.S.

## SUPPLEMENTARY FIGURES

**Supplementary Figure 1:**
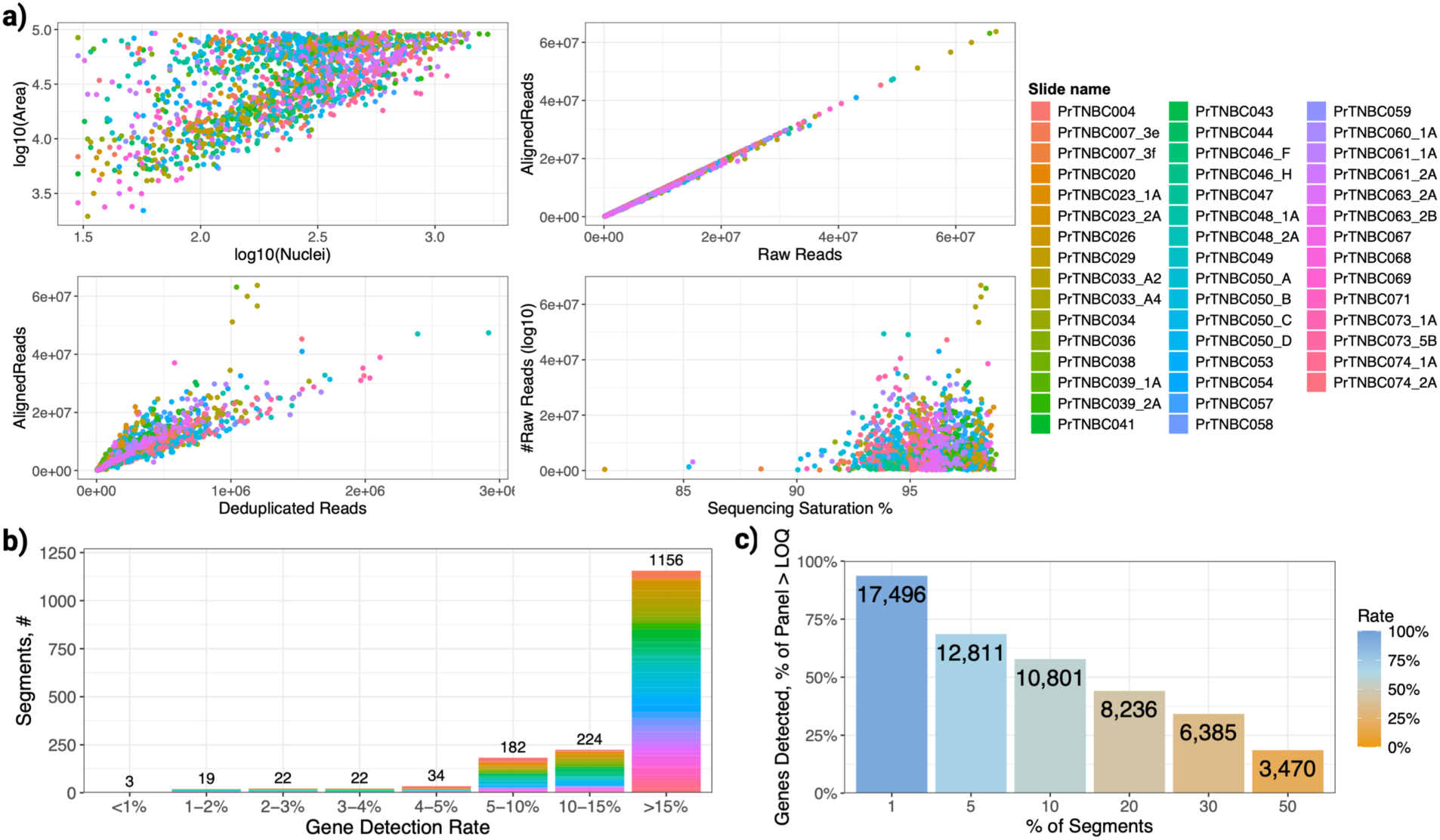
Quality control plots. **a)** Plots of area, # nuclei, raw reads, aligned reads, deduplicated reads, and sequencing saturation after filtering AOIs with the following quality control cutoffs: *minAOIReads = 1000, percentTrimmed = 80, percentStitched = 80, percentAligned = 80, percentSaturation = 50, minNegativeCount = 1, maxNTCCount = 5000, minNuclei = 30, minArea = 1000, LOQ = 2*. **b)** Plot of gene detection rate across segments colored by slide. Only AOIs expressing > 5% of genes were maintained (1562 AOIs). **c)** Plots of genes detected with LOQ > 2 across all AOIs/segments. Only genes expressed in >10% of AOIs were maintained (10801 genes).

**Supplementary Figure 2:**
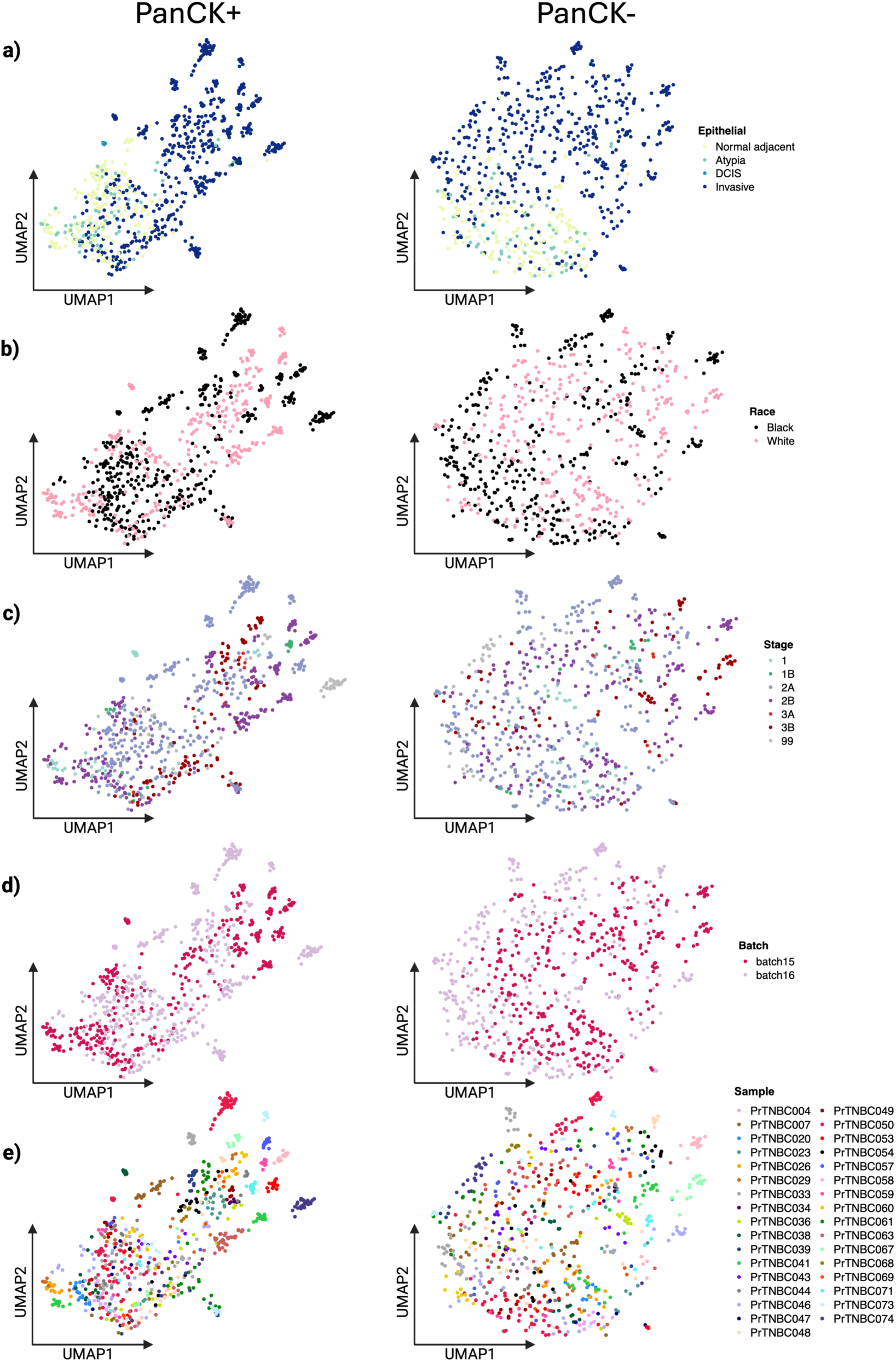
UMAPs across metadata. UMAP plots across PanCK+ and PanCK- AOIs, colored by **a)** epithelial type, **b)** race, **c)** stage, **d)** batch, and **e)** sample. The batch UMAPs (**S2d**) show that there is no batch effect.

**Supplementary Figure 3:**
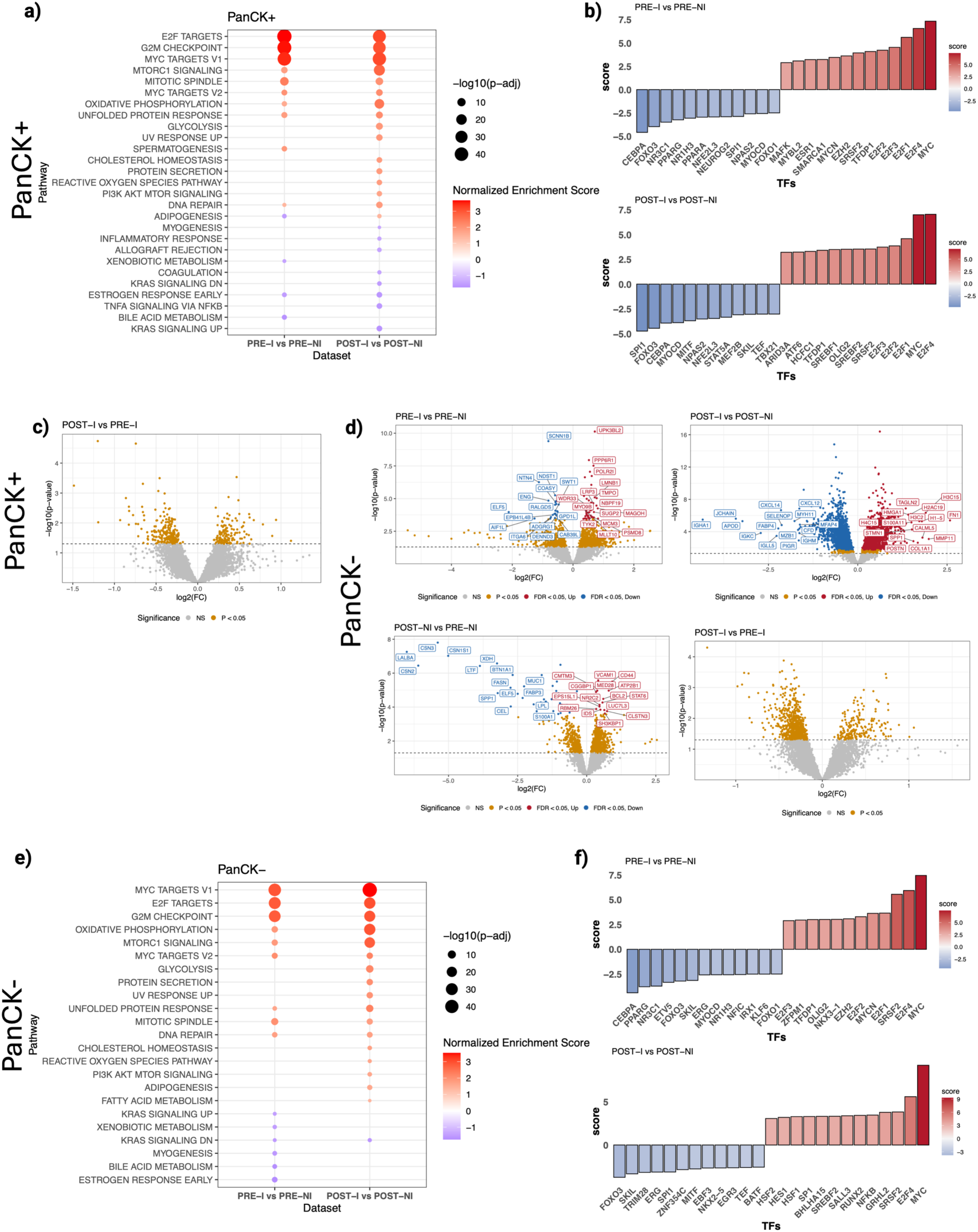
Supplementary plots for the characterization of epithelia and TME. **a)** Enrichment of the PRE-I vs PRE-NI and POST-I vs POST-NI comparisons in PanCK+ AOIs (Epithelia). **b)** Transcription factor (TF) target analysis of the PRE-I vs PRE-NI and POST-I vs POST-NI comparisons in PanCK+ AOIs (Epithelia). **c)** Volcano plot of the POST-I vs PRE-I comparison in PanCK+ AOIs shows that there is no statistically significant difference (all FDR > 0.05) when directly comparing invasive epithelia in PRE vs POST. **d)** Volcano plots for the PRE-I vs PRE-NI, POST-I vs POST-NI, POST-NI vs PRE-NI, and POST-I vs PRE-I comparisons in PanCK- AOIs (TME). **e)** Enrichment of the PRE-I vs PRE-NI and POST-I vs POST-NI comparisons in PanCK- AOIs (TME). **f)** TF target analysis of the PRE-I vs PRE-NI and POST-I vs POST-NI comparisons in PanCK- AOIs (TME).

**Supplementary Figure 4:**
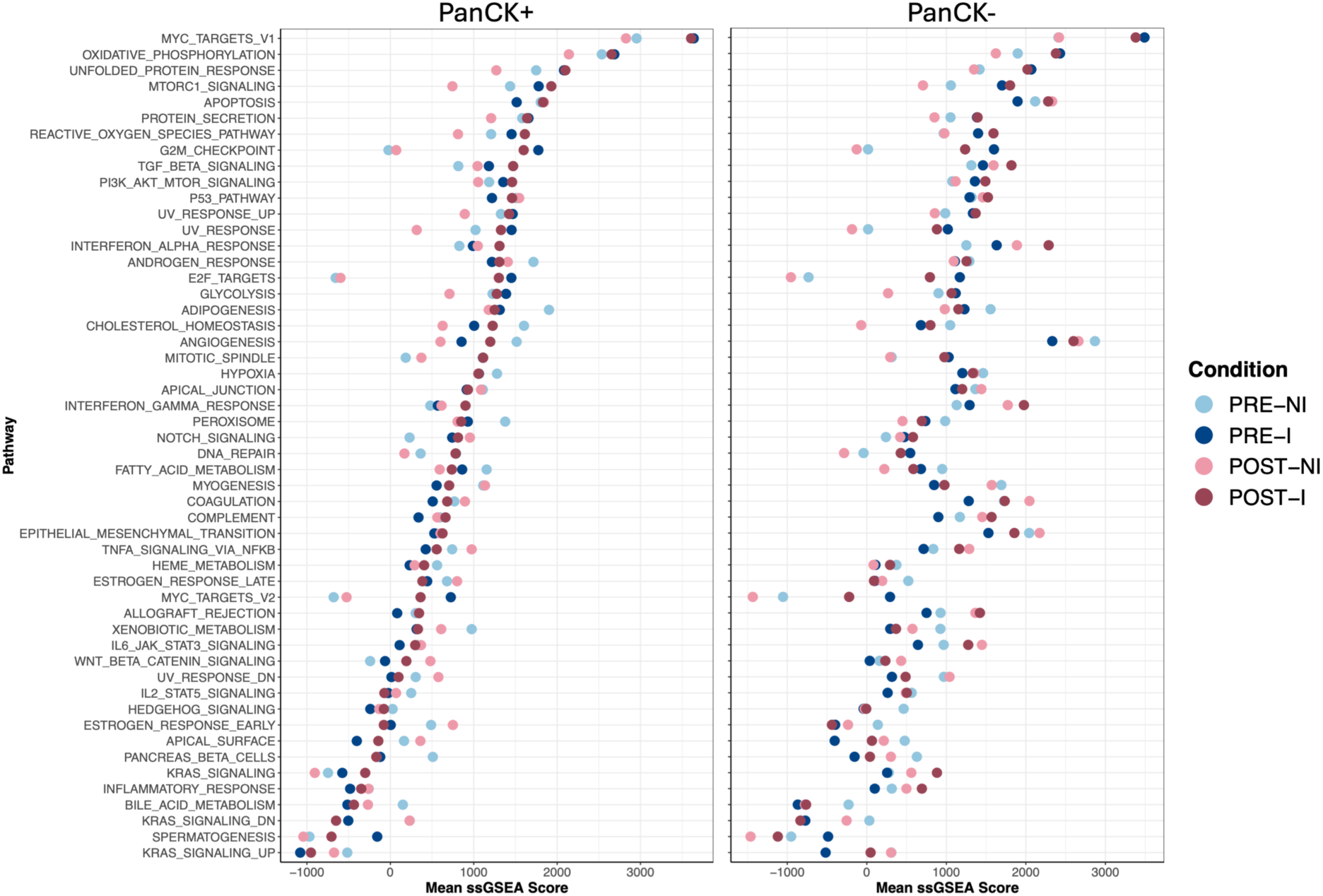
Average ssGSEA score across groups. **a)** Single sample gene set enrichment (ssGSEA) was performed for each AOI using MSigDB Hallmark pathways. PanCK+ AOIs were grouped into PRE-NI, PRE-I, POST-NI, and POST-I and the average ssGSEA score in each group for each pathway was plotted. **b)** Same as a), but for PanCK- AOIs. This provides supporting evidence for the differential expression-based enrichment results.

**Supplementary Figure 5:**
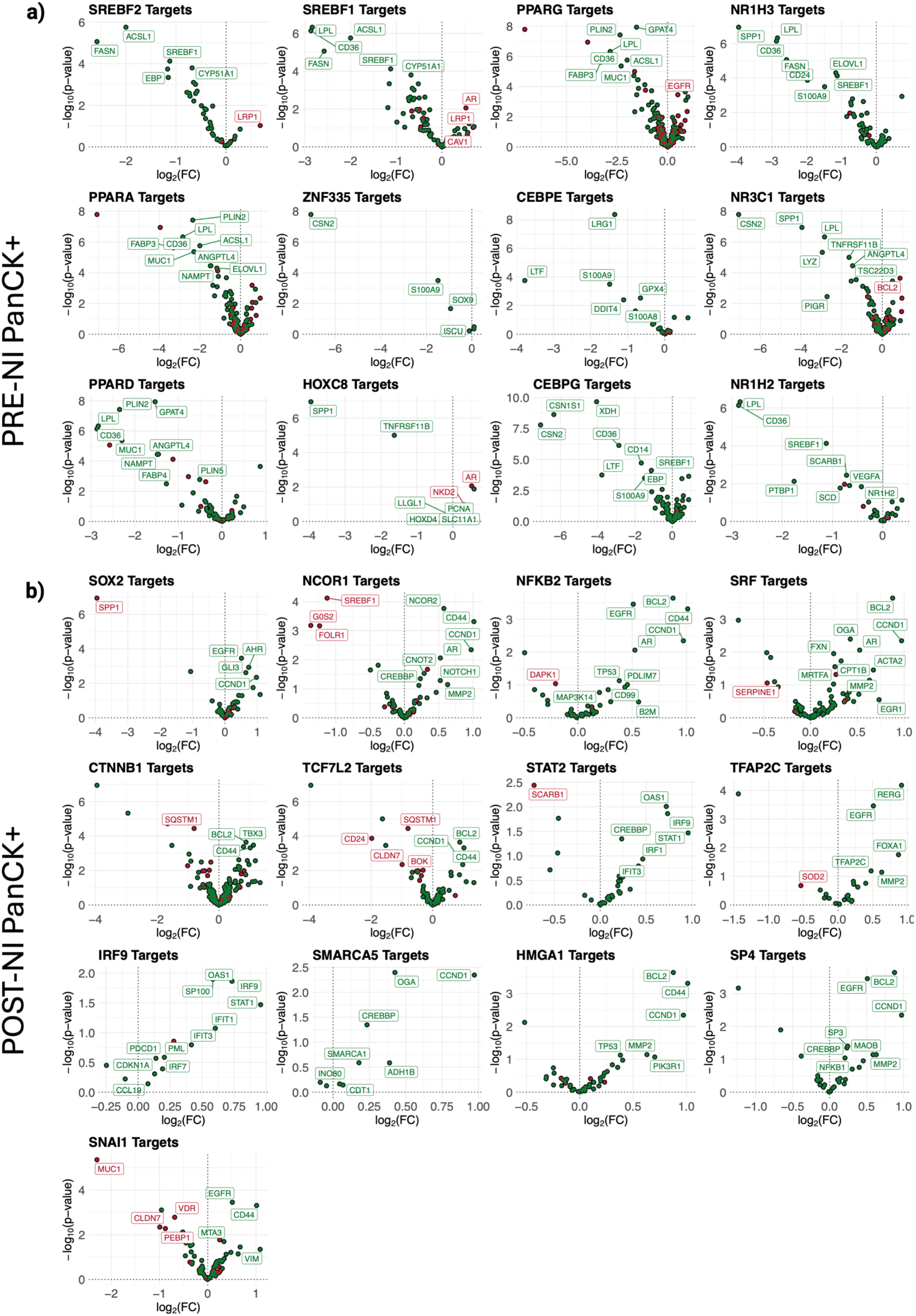
Downstream target genes driving the differential TF activity in POST-NI vs PRE-NI PanCK+ AOIs (Epithelia). **a)** Log2FC and p-values of downstream genes of the most significant TFs in PRE-NI PanCK+ AOIs. Green indicates genes that are activated by the TF and red indicates genes suppressed by the TF. Positive log2FC indicates that the gene is upregulated in POST-NI, while a negative log2FC indicates that the gene is upregulated in PRE-NI. **b)** Log2FC and p-values of downstream genes of the most significant TFs in POST-NI PanCK+ AOIs.

**Supplementary Figure 6:**
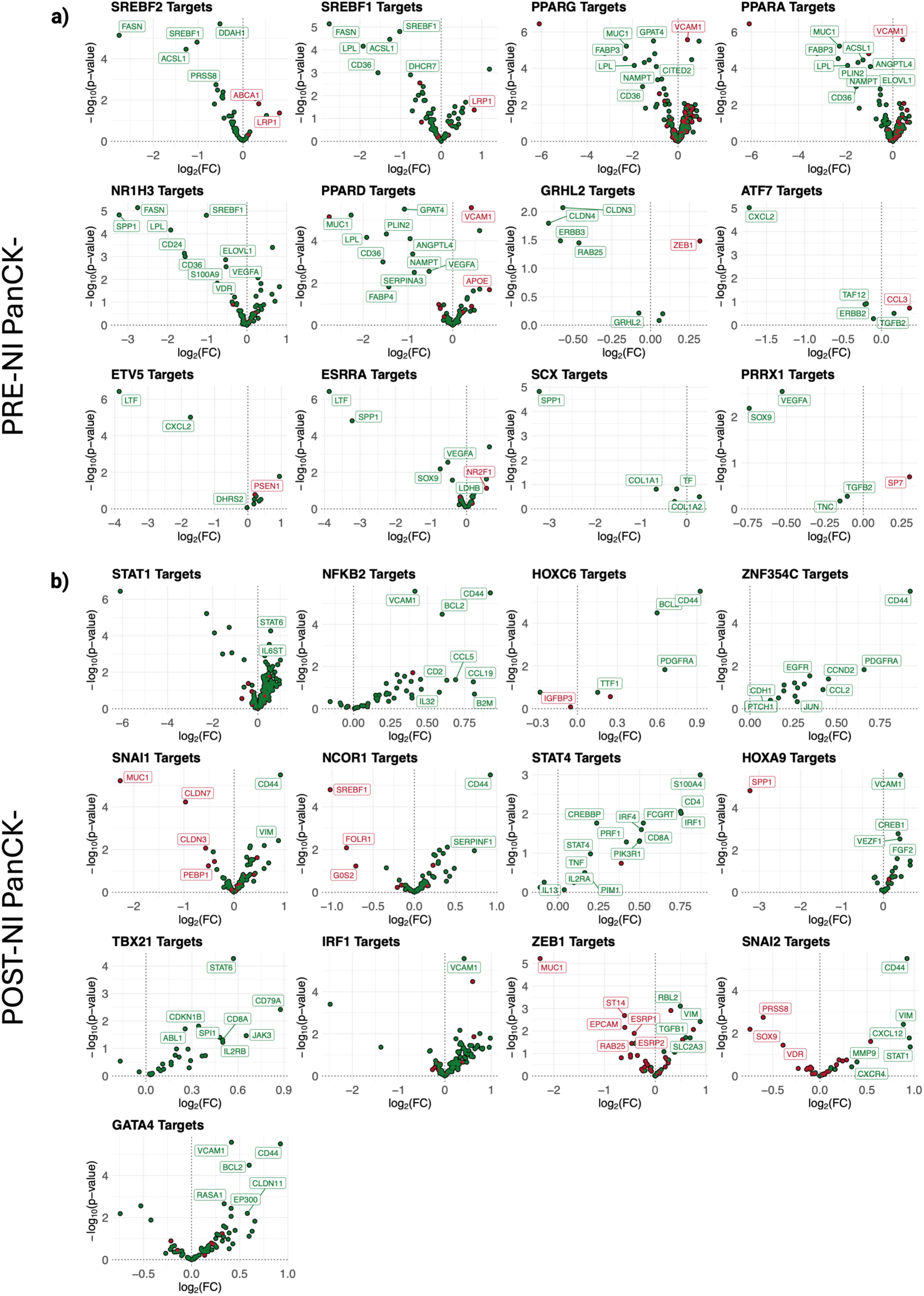
Downstream target genes driving the differential TF activity in POST-NI vs PRE-NI PanCK- AOIs (TME). **a)** Log2FC and p-values of downstream genes of the most significant TFs in PRE-NI PanCK- AOIs. Green indicates genes that are activated by the TF and red indicates genes suppressed by the TF. Positive log2FC indicates that the gene is upregulated in POST-NI, while a negative log2FC indicates that the gene is upregulated in PRE-NI. **b)** Log2FC and p-values of downstream genes of the most significant TFs in POST-NI PanCK- AOIs.

**Supplementary Figure 7:**
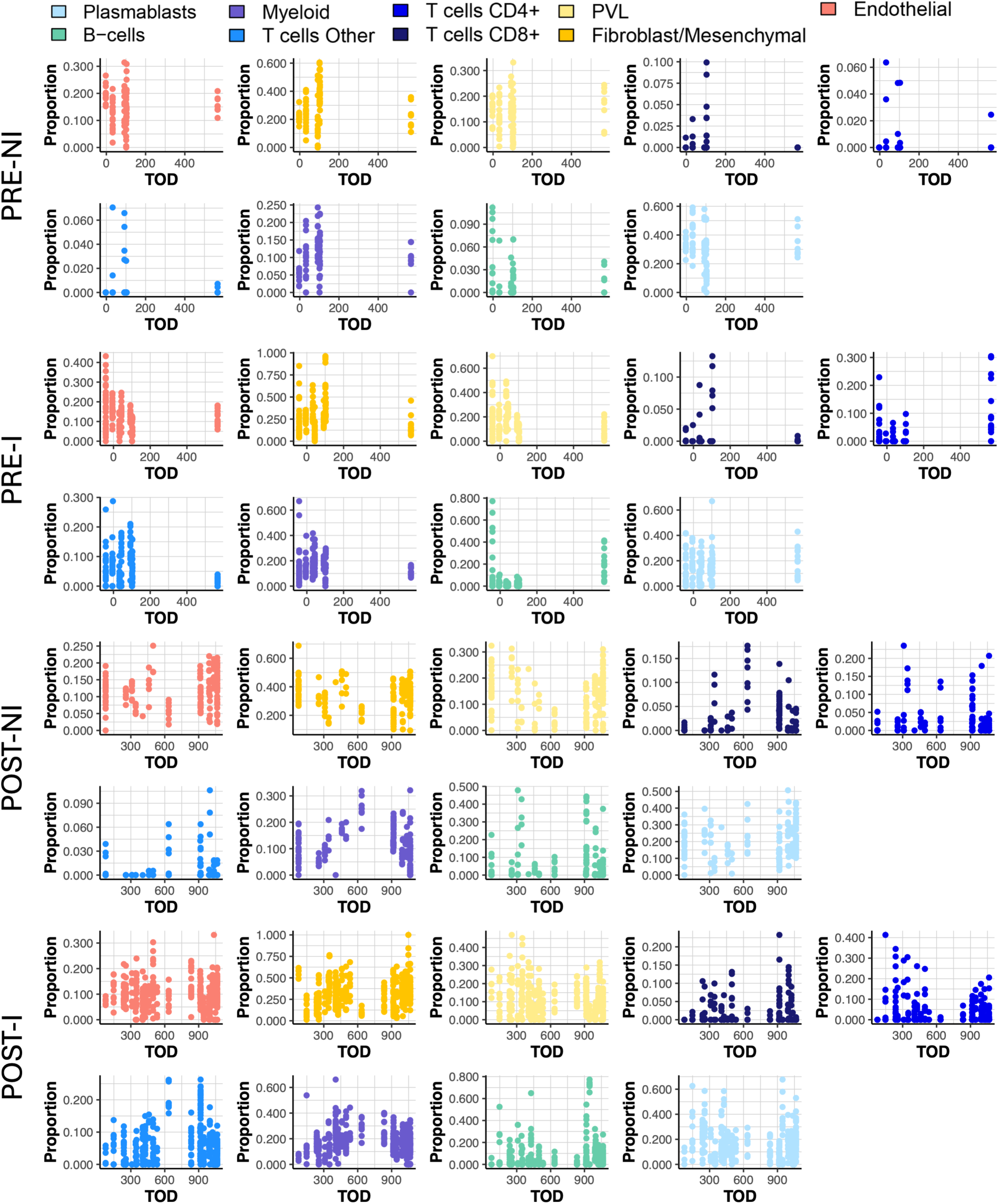
Distribution of cell type proportions arranged by time of diagnosis (TOD). Distribution of the calculated proportions (by SpatialDecon^37^) of each cell type in each ROI, arranged by time of diagnosis and separated by group: PRE-NI, PRE-I, POST-NI, and POST-I.

**Supplementary Figure 8:**
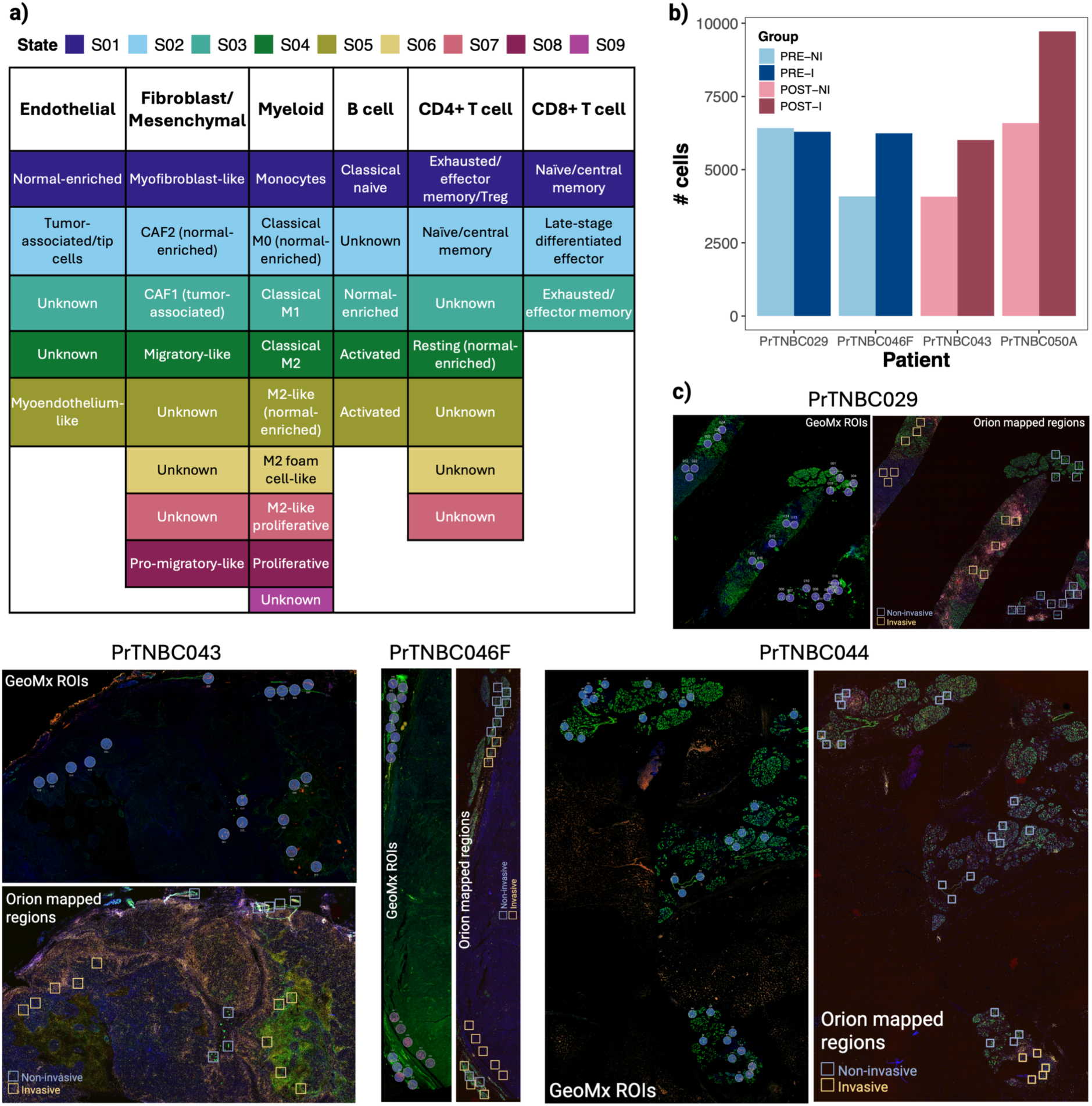
Table of cell state labels from Ecotyper and region mapping on slides used for Orion. **a)** Table of all cell states associated with each cell type in Ecotyper, taken from Table S4 of the original publication^41^. **b)** Total # cells coming from all square regions considered for Orion analysis from each slide, across non-invasive and invasive regions. PrTNBC029/ PrTNBC046F are PRE and PrTNBC043/PrTNBC050A are POST. **c)** Slide annotations show how 300um square regions were selected on slides used for Orion to match the locations of ROIs selected for GeoMx spatial sequencing. PrTNBC044 (POST diagnosed <1 year after delivery) was added comparing CD3+ and PanCK+ cell counts across pseudotime, along with PrTNBC043 (POST diagnosed 1-2 years after delivery) and PrTNBC050A (POST diagnosed 2-3 years after delivery, slide image in Figure 4c).

**Supplementary Figure 9:**
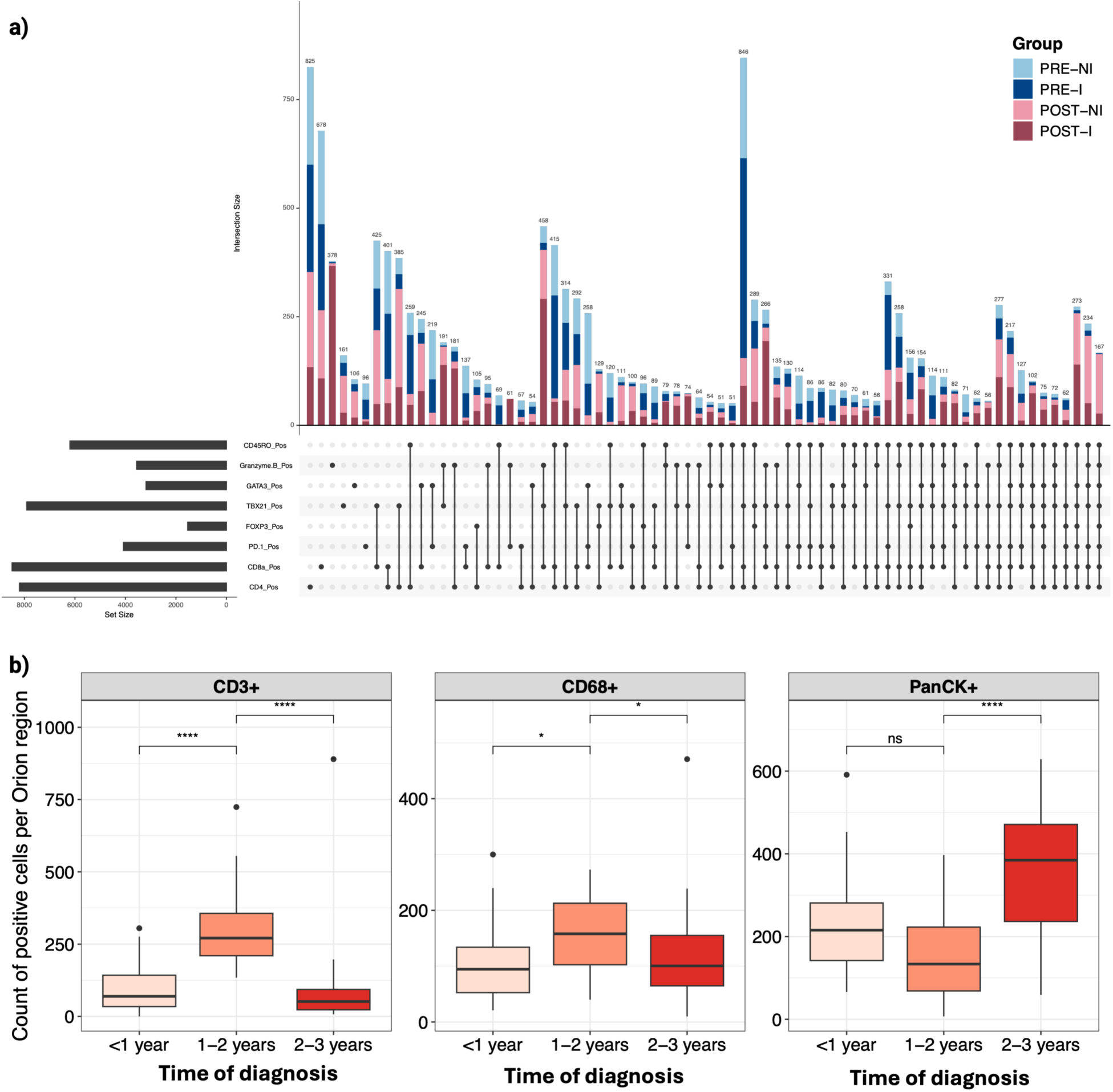
Orion supplementary figures. **a)** Distribution of all T cell states identified by unique marker combinations (with >50 representative cells) across all CD3+ cells, colored by group. 2 PRE and 2 POST slides were used for Orion whole slide multiplexed imaging (see Supplementary Figure 8c). **b)** Boxplots of CD3+ (T cell), CD68+ (Macrophage), and PanCK+ (Epithelial cell) counts across all regions considered in Orion staining. As shown in Supplementary Figure 8c, Orion regions were selected to match as closely as possible to GeoMx ROIs. 1 representative sample was chosen for each pseudo-timepoint. The woman diagnosed 1-2 years after delivery showed significantly more T cells and CD68+ macrophages per ROI than the other time points, in line with the transcriptomic findings.

**Supplementary Figure 10:**
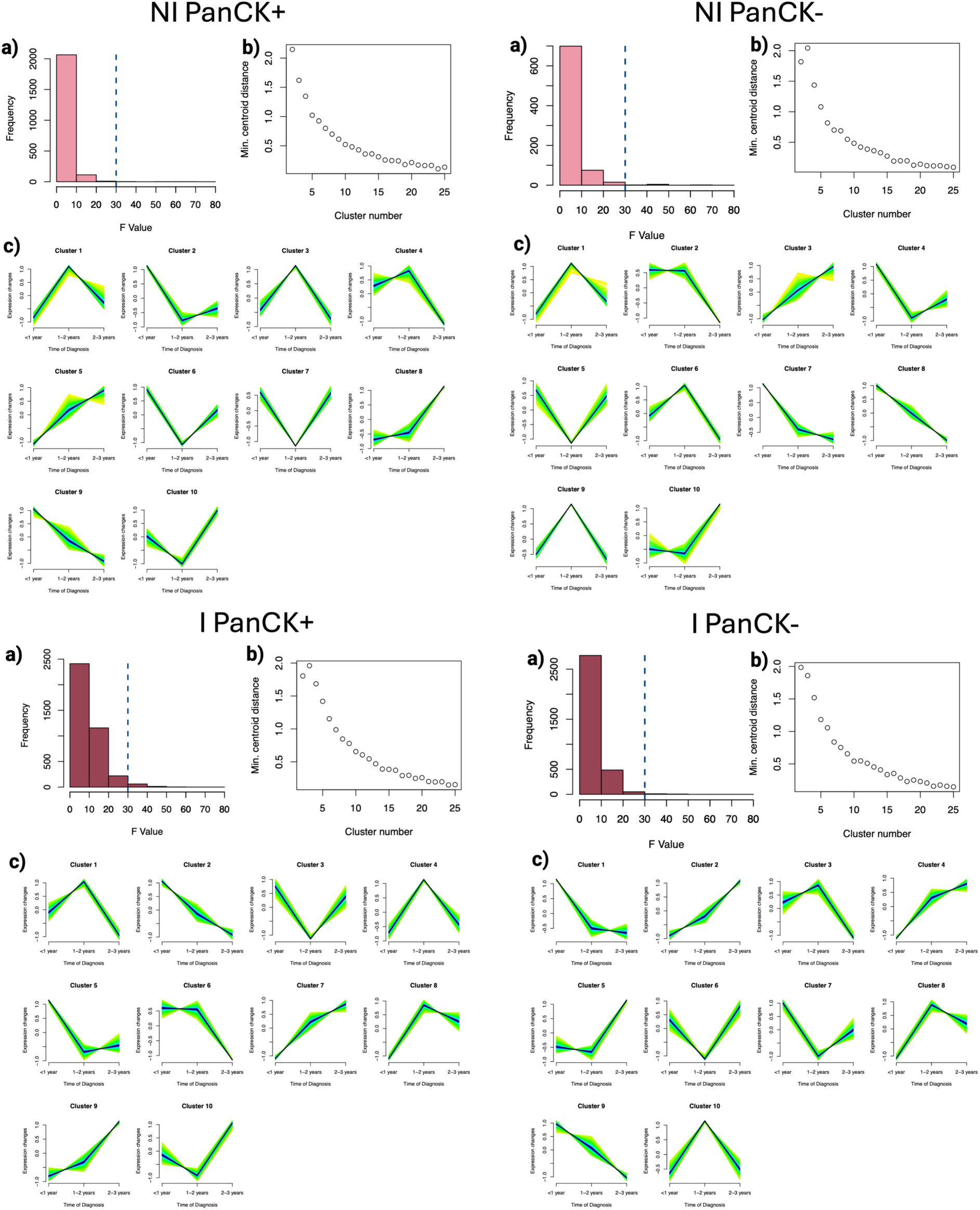
Pseudotime analysis plots. Post-SPLS plots, separated by group (POST-NI PanCK+, POST-NI PanCK-, POST-I PanCK+, and POST-I PanCK-). In each: **a)** Plots of ANOVA F-value for all genes identified by SPLS as important for separating between the three timepoints. F-values > 30 were removed for downstream analysis. **b)** Plots of minimum centroid distance for varying numbers of clusters prior to performing Mfuzz. 10 clusters were chosen for all groups. **c)** Gene clusters identified by Mfuzz for each group. Clusters with similar trends (increasing across time, peaking in timepoint 2, etc) were combined for further functional analysis.

**Supplementary Figure 11:**
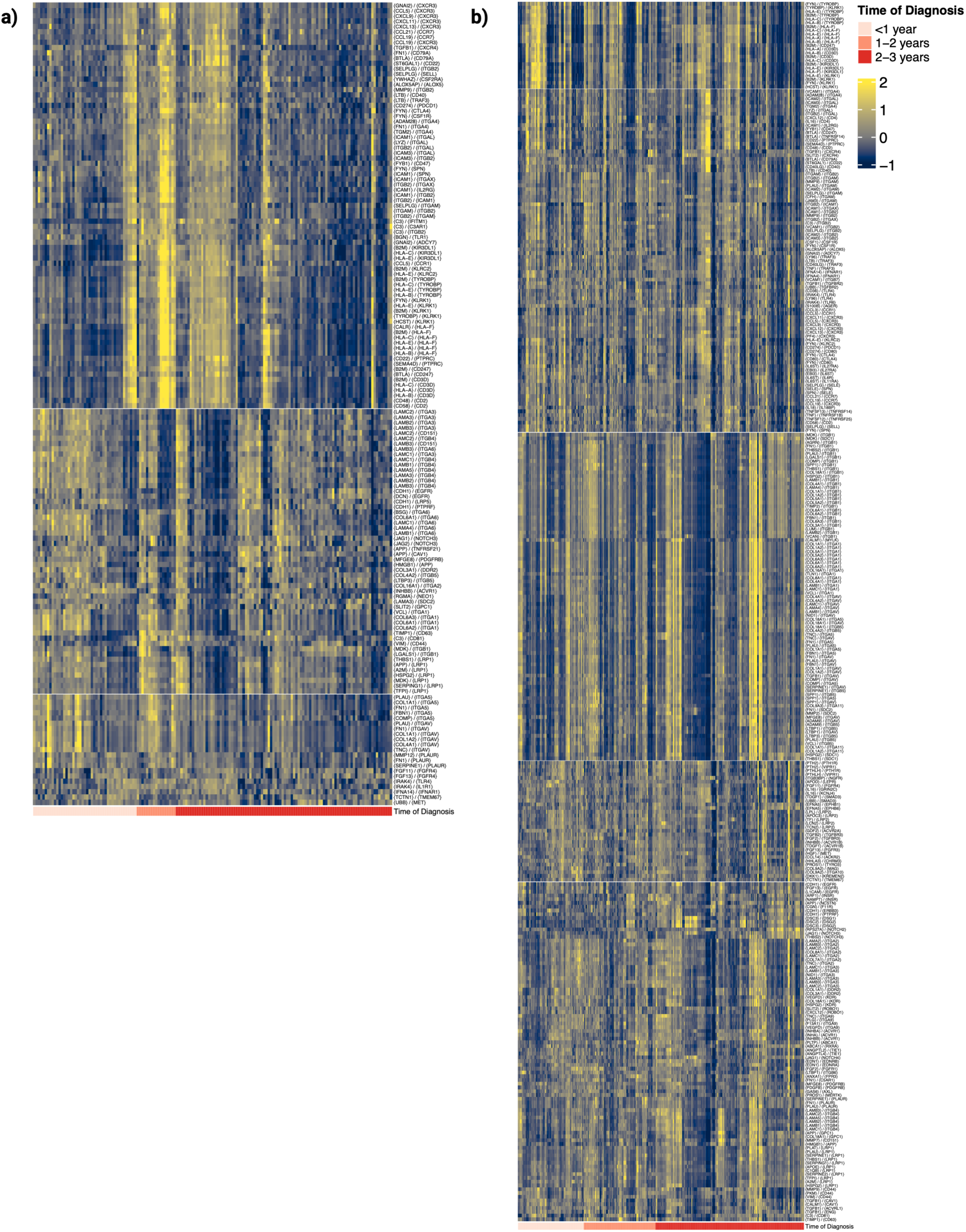
Heatmaps of Ligand-Receptor interactions in POST-NI and POST-I TME. **a)** Heatmap of all ligand-receptor ({L} / {R}) interactions identified by BulkSignalR in POST-NI (non-invasive) PanCK- AOIs, arranged from earliest to latest time of diagnosis after delivery. Heatmap values are gene signature scores for each interaction, calculated by a weighted sum involving the average of L and R gene z-scores (half of the score) and the average of the target gene z-scores (other half of the score)^43^. AOIs from women diagnosed 1-2 years post-involution and the earliest AOIs from women diagnosed 2-3 years post-involution (up to 480 days) show a distinct high score for a set of 87 LR interactions; the top 50 of these are shown in Figure 5c. **b)** Heatmap of all L-R interactions identified by BulkSignalR in POST-I (invasive) PanCK- AOIs, arranged from earliest to latest time of diagnosis. There are no clear trends in LR interactions across pseudotime in the invasive TME as there are in the non-invasive TME (Supplementary Figure 11a).

